# Detecting directed motion and confinement in single-particle trajectories using hidden variables

**DOI:** 10.1101/2024.04.18.589798

**Authors:** François Simon, Guillaume Ramadier, Inès Fonquernie, Janka Zsok, Sergiy Patskovsky, Michel Meunier, Caroline Boudoux, Elisa Dultz, Lucien E. Weiss

## Abstract

Single-particle tracking is a powerful tool for understanding protein dynamics and characterizing microenvironments. As the motion of unconstrained nanoscale particles is governed by Brownian diffusion, deviations from this behavior are biophysically insightful. However, the stochastic nature of particle movement and the presence of localization error pose a challenge for the robust classification of non-Brownian motion. Here, we present *aTrack*, a versatile tool for classifying track behaviors and extracting key parameters for particles undergoing Brownian, confined, or directed motion. Our tool quickly and accurately estimates motion parameters from individual tracks.

Further, our tool can analyze populations of tracks and determine the most likely number of motion states. We show the working range of our approach on simulated tracks and demonstrate its application for characterizing particle motion in cells and for biosensing applications. aTrack is implemented as a stand-alone software, making it simple to analyze track data.

## Main

Single-particle tracking (SPT) is a valuable tool for characterizing protein activity (1–3). While Brownian diffusion is commonly observed inside of cells, deviations from this behavior are of major interest and can provide key biophysical insight (4–6). Classically, non-Brownian behaviors have been identified by fitting a power law to the mean-squared displacements (MSD) as a function of the time lag *τ*, (MSD = Γ·*τ*^*α*^) (6), where subdiffusive motion has an *α* exponent < 1 and superdiffusion has an exponent > 1. While fitting the MSD curve can therefore be used for categorizing motion, the anomalous exponent and the generalized diffusion coefficient provide little insight into the combination of underlying biological forces, e.g., diffusion, directed motion, and confinement.

To accurately model the motion encountered in cellular contexts, physics-based models have been developed to explicitly describe interactions that give rise to nonlinear MSDs. For example, superdiffusion can arise from the linear movement powered by molecular motors (7), and subdiffusive motion can arise from confinement (8, 9). In such models, the observed track positions result from a stochastic process described by hidden (unobserved) variables that evolve with time.

In the context of random processes, Maximum Likelihood Estimation (MLE) is used to determine the underlying parameters of the model. MLE consists in computing the probability of observing a track given the model and its parameters. The main challenge in models with hidden variables is that they require computing the integral of a joint probability density over all possible hidden variables. The few methods that use this type of models perform this integration step sampling methods that are quite slow and inaccurate (8, 9). Thus, designing a hidden-variable method that benefits from accurate and efficient integration is an important challenge for improving the reliably and ease-of-use of the physics-based models. Further, when the underlying physical model is undecided, statistical tests can be applied to identify the most appropriate model (10–12).

Current hidden-variable models applied to non-Brownian diffusion have three disadvantages: 1) They perform the integration step using either coarse-grained approximations (7) or sampling method making them either inaccurate or slow; 2) they are specialized in either confinement or directed motion but do not treat both types of motion; 3) they typically do not allow variations of the potential well position or the speed of the directed motion, properties that are frequently encountered in biological systems.

Here, we present aTrack, an analysis tool that alleviates the limitations mentioned above. Our approach uses a versatile motion model that considers the relationships between the observed track, the real particle positions (localization error and Brownian motion), as well as the influence of a non-Brownian variable that can either be the potential well for confined diffusion or the velocity vector for directed motion. The main innovations of this model are: 1) it uses analytical recurrence formulas to perform the integration step for complex motion, improving speed and accuracy; 2) it handles both confined and directed motion; 3) anomalous parameters, such as the center of the potential well and the velocity vector are allowed to change through time to better represent tracks with changing directed motion or confinement area; and lastly 4) for a given track or set of tracks, aTrack can determine whether tracks can be statistically categorized as confined or directed, and the parameters that best describe their behavior, for example, diffusion coefficient, radius of confinement, and speed of directed motion.

We validate the approach on simulated data and demonstrate its versatility for analyzing a variety of experimental SPT data, including particle diffusion in an optical trap, detection of motile bacteria with gold nanoparticles, and motion characterization of spindle pole bodies in budding yeast.

## Results

### Accounting for hidden variables that characterize confined and persistent motion

#### Modeling non-Brownian motion

We model noisy tracks undergoing confined, Brownian, and directed motion by considering four relations at each time step: (1) there is a Brownian diffusion step followed by (2) an anomalous step (Fig. 1a-b); (3) the hidden anomalous variable, *h*, can evolve according to a Gaussian distribution; and (4) localization error is incorporated as a Gaussian-distributed noise term added to the underlying real position to produce the observed positions. This model encompasses a variety of motion types depending on the model parameters, as illustrated in Fig. 1c. For example, particles can be immobile, where the observed displacements are only due to localization error; tracks can undergo Brownian motion, as well as anomalous super- and subdiffusion; or multiple motion mechanisms can also occur simultaneously, (e.g. diffusion and drift, or changes in directed motion direction and speed). The model can also account for confinement in an area of variable radius (8, 9, 13) as well as a diffusing potential well. In our model, the velocity is the characteristic parameter of directed motion and the confinement factor represents the force within a potential well. More precisely, the confinement factor *l* is defined such that at each time step the particle position is updated by *l* times the distance particle/potential well center (see the Methods section for more details).

**Figure 1.**
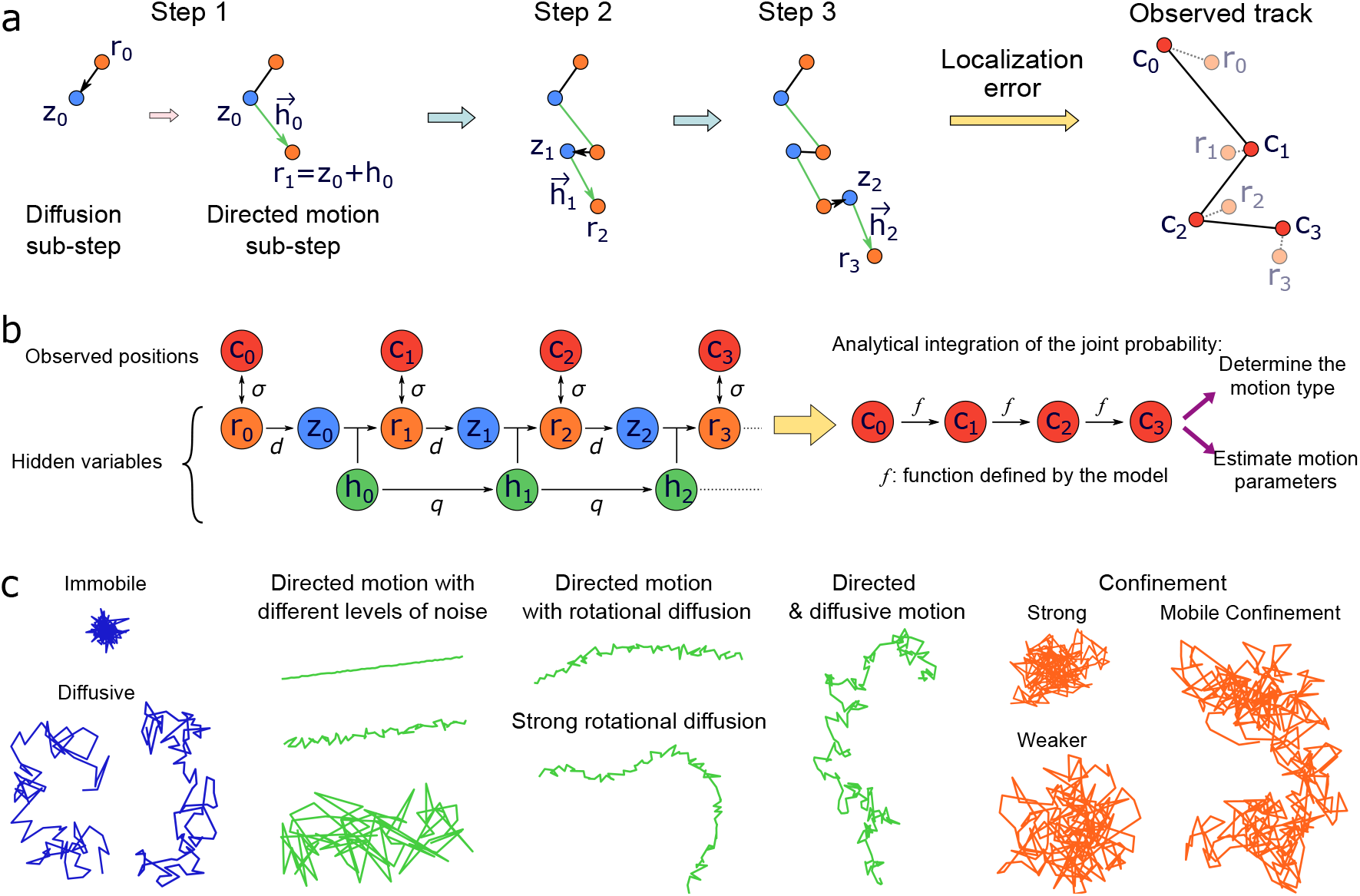
Principle of *aTrack*. **a**: The track-generation steps with our motion model shown here for directed motion. Each time step is decomposed into sub-steps, namely diffusion and anomalous motion (either directed or confining) with *r*_*i*_ the real positions, *z*_*i*_ an intermediate position, *h*_*i*_ the anomalous variable, and the generation of observed (measured) positions, *c*_*i*_. The variables *c*_*i*_ −*r*_*i*_ and *r*_*i*+1_ −*r*_*i*_ follow Gaussian distributions with mean 0 and standard deviations *σ* and *d*, respectively. The sub-step from *z*_*i*_ to *r*_*i*+1_ is deterministic and the anomalous variable *h*_*i*_ can also evolve with a standard deviation *q*. **b**: Graph representation of aTrack showing the motion model (*left*), analytical integration (*middle*), and outputs (*right*). To compute the track probability, we integrate over the hidden variables. This results in an analytical recurrence formula that is used to determine the type of motion and to estimate the parameters of the motion. **c**: Examples of tracks that can be produced with our motion model.

#### Determining the type of motion

To categorize the type of motion from a measured trajectory, we first calculate the likelihood that a track belongs to each considered motion class (diffusion, directed, or confinement) and then perform a statistical comparison between the likelihoods. To do the former, we must integrate the joint probability density function of a track for all the modeled hidden variables, e.g. all the real positions and the potential well positions or the velocity vectors. Since our model is a multivariate Gaussian process expressed as the product of univariate Gaussian functions, we can perform the integration step using analytical recurrence formulas (see Supplementary information) (14, 15). The recurrence formula enables our model computation time to scale linearly with the number of time points.

We can apply these formulas to determine the best sets of parameters and the maximum likelihoods, then use the ratio between the maximum likelihood assuming Brownian diffusion (null hypothesis) and the maximum likelihood assuming confinement or directed motion as the alternative hypothesis to build a likelihood ratio test (16, 17). Fig. S2 shows that these likelihood ratios, *ρ* = *l*_Brownian_/*l*_confined_ or *ρ* = *l*_Brownian_/*l*_directed_ are systematically skewed towards 1 when particles follow Brownian diffusion. Conversely, *ρ* is skewed toward zero when applying the directed-diffusion test to directed tracks or the confined-diffusion test to confined tracks. These properties enable the likelihood ratio to serve as a robust proxy for the p-value as it overestimates it. As a consequence, if *ρ* < *X* with X the type I error rate (e.g. 0.05), we can reject the null hypothesis with a confidence of 1 − *X*. Relying on the skewness of the likelihood ratio to obtain an upper bound of the p-value is a simple way to categorize the type of motion, but sometimes a more sensitive test is needed. In such cases, one can use simulations to better estimate the p-value (See Supplemental Information: Statistical test for more details).

To estimate the impact of the track length on the classification, we simulated tracks either in confined diffusion or in linear motion (without diffusion) of varying lengths (5-400 steps for confined, 4-100 steps for directed) and computed the likelihood ratios. As expected, the classification certainty increases with the track length; where the inverse of log *ρ* increases with the number of steps (Fig 2a). Of course, the statistical certainty depends on the track parameters (Fig 2b). In the range of evaluated anomalous parameters, the significance of the test increased with the confinement factor and the directed motion velocity. To determine the useful range more systematically, we varied the track length and either the confinement factor for confined motion or the velocity for directed motion, and computed the average likelihood ratios (Fig 2c). A low average ratio indicates significantly low p-values for most tracks as this ratio is an overestimate of the p-value. We see that the test is significant as long as the anomalous parameter is high enough or the track length is high enough. Note that increasing the confinement factor so much that the confinement radius becomes similar or smaller than the localization error will impair the capacity of the test.

**Figure 2.**
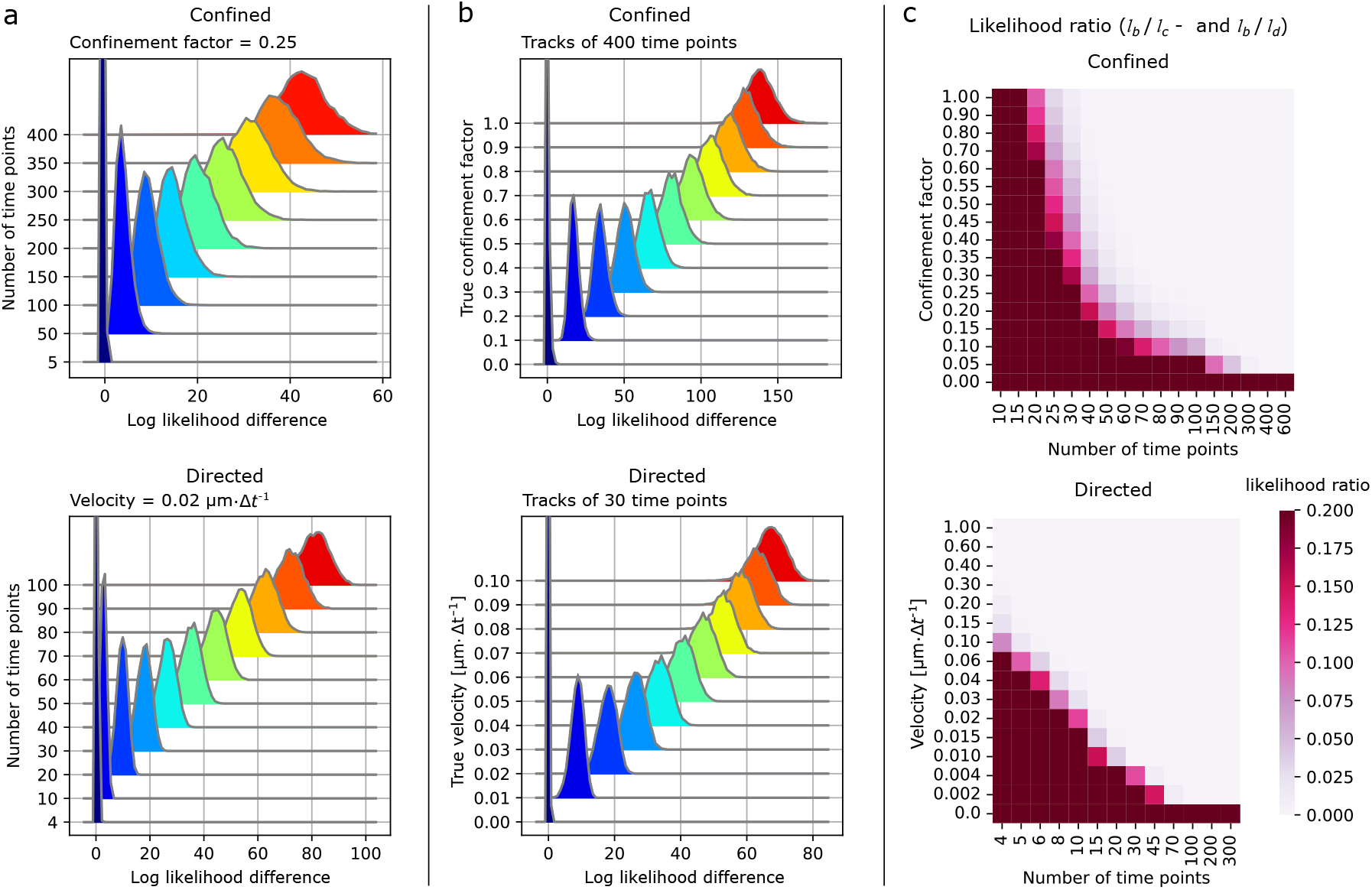
Determining the motion type using a likelihood-ratio test. **a-b**: Probability distributions of the difference between the log of the maximum likelihood of the alternative hypothesis (either confinement *L*_*c*_ or directed *L*_*d*_ ) and the null hypothesis (Brownian diffusion *L*_*b*_) for single tracks (10,000 tracks). Confinement factor *l* = 0.25 and velocity *v* = 0.02 µm·Δt^−1^. **a**: Effect of the number of time points in a track on its log difference (*L*_*c*_ − *L*_*b*_ for confined tracks) and (*L*_*d*_ − *L*_*b*_ for directed tracks). **b**: The ability to distinguish confinement and directed motion from diffusion as a function of the confinement factor and particle velocity, respectively. **c**: heatmaps of the likelihood ratios *l*_*b*_/*l*_*c*_ (confined) or *l*_*b*_/*l*_*d*_ (diffusive) varying both the anomalous diffusion parameter and the track length. Mean of 10,000 tracks. **a-c**: When not stated otherwise, the track parameters were as following. Localization error *σ* = 0.02 µm. Confined tracks: diffusion length *d* = 0.1 µm. Directed tracks: *d* = 0.0 µm (no diffusion), constant speed and orientation.

#### Characterizing confinement

To characterize confined trajectories, aTrack estimates several parameters, namely the diffusion coefficient *D* and the diffusion length 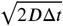 where Δ*t* is the single-frame time step; the confinement factor *l*, which is proportional to the spring constant of the potential well; the diffusion coefficient of the confinement area *D*_*c*_; and the localization error *σ*; and The confinement radius, which is proportional to 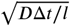, can also be calculated (See the methods section for more details).

To measure the precision of parameter estimation, we simulated tracks with different confinement factors, *l*, (Fig. 3a). We then used aTrack to estimate the parameters for each track of 200 time steps (Fig. 3b). The average diffusion coefficient, confinement factor, and confinement radius were accurate over the range of confinement factors, 0 to 1 per time step. Panel Fig. 3c shows the working range for calculating the confinement factor as a function of track length and confinement factor. Longer tracks result in better parameter estimation. Similarly, our method correctly estimates the confinement radius (Fig. S3d). In a second set of simulations, we tested the reliability of our predictions when the confinement area is moving (Fig. 3d, Fig. S3e).

**Figure 3.**
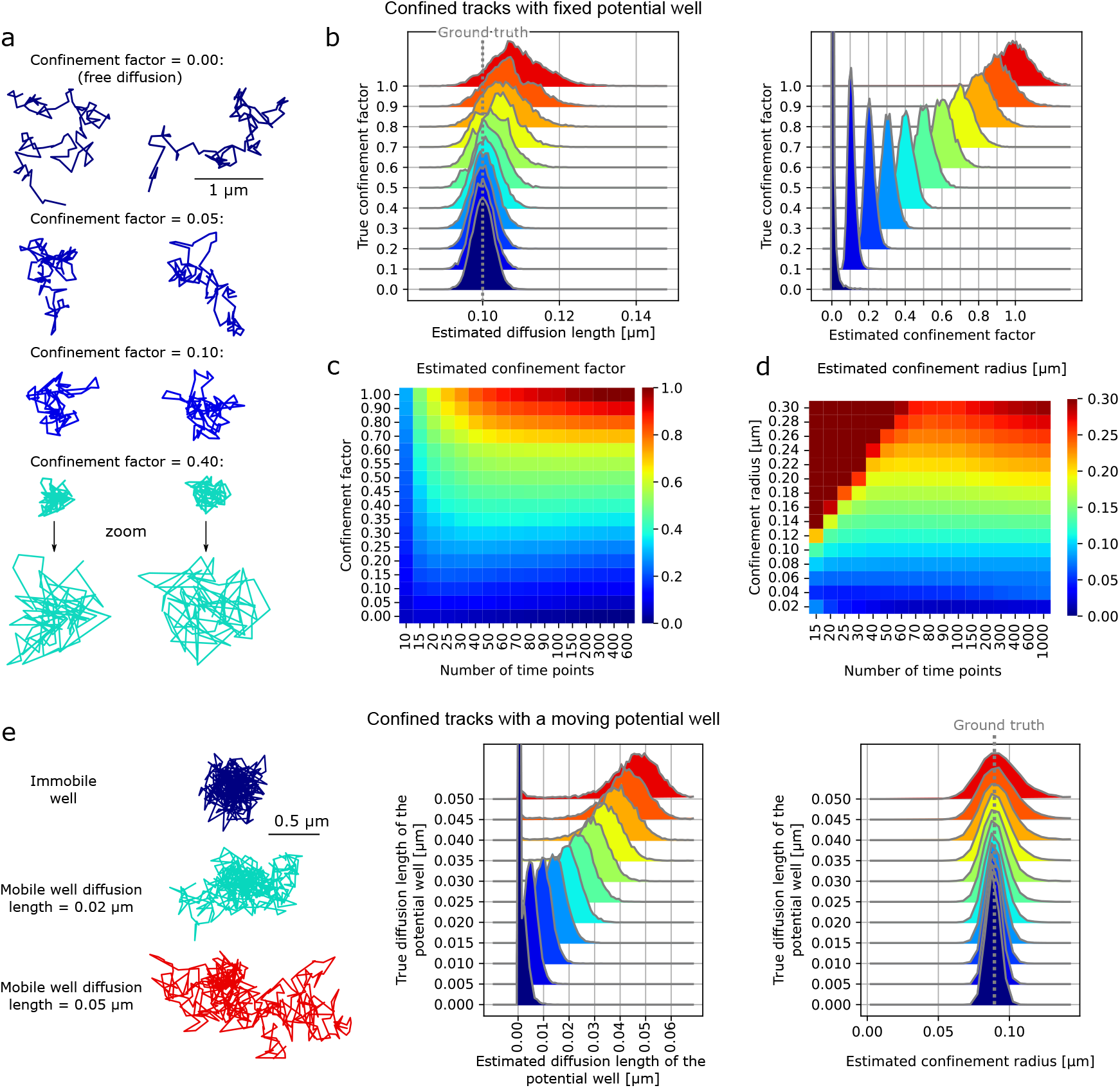
Characterizing confinement with *aTrack*. **a-d**: Confinement of tracks with a fixed potential well. **a**: Examples of simulated tracks with different confinement factors. **b**: Histograms of the estimated parameters for individual tracks of 200 time points varying the number of time points in tracks. **c-d**: Heatmaps of the mean estimated confinement factor and confinement radius depending on the track length and the confinement factor (per time step) or radius respectively. **e**: Confinement of tracks with a moving potential well (Brownian motion). Left: simulated tracks with different diffusion length of the potential well. Right: histograms of the estimated diffusion length of the the potential well and confinement radius 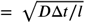 varying the actual diffusion length of the potential well. Confinement factor = 0.1 per time step. **a-d**: 10,000 tracks per condition. *d* = 0.1 µm, Localization error *σ* = 0.02 µm. See Fig. S3 for complementary results.

#### Characterizing directed motion

Superdiffusive behavior, in particular directed motion, occurs in a variety of circumstances, such as molecular motor-mediated active transport (7, 18) and polymerase processivity (19). Localization error complicates velocity estimation, especially when the localization error is relatively large. We simulated noisy tracks undergoing linear motion at various speeds (Fig. 4a) with a localization error of 20 nm·Δ*t*^−1^ and estimated the velocity per tracks. Fig. 4b shows the velocity estimates for tracks with 30 time points. By varying both the track length and the velocity, we found our method to be reliable for a wide range of velocities as long as tracks were long enough (Fig. 4c).

**Figure 4.**
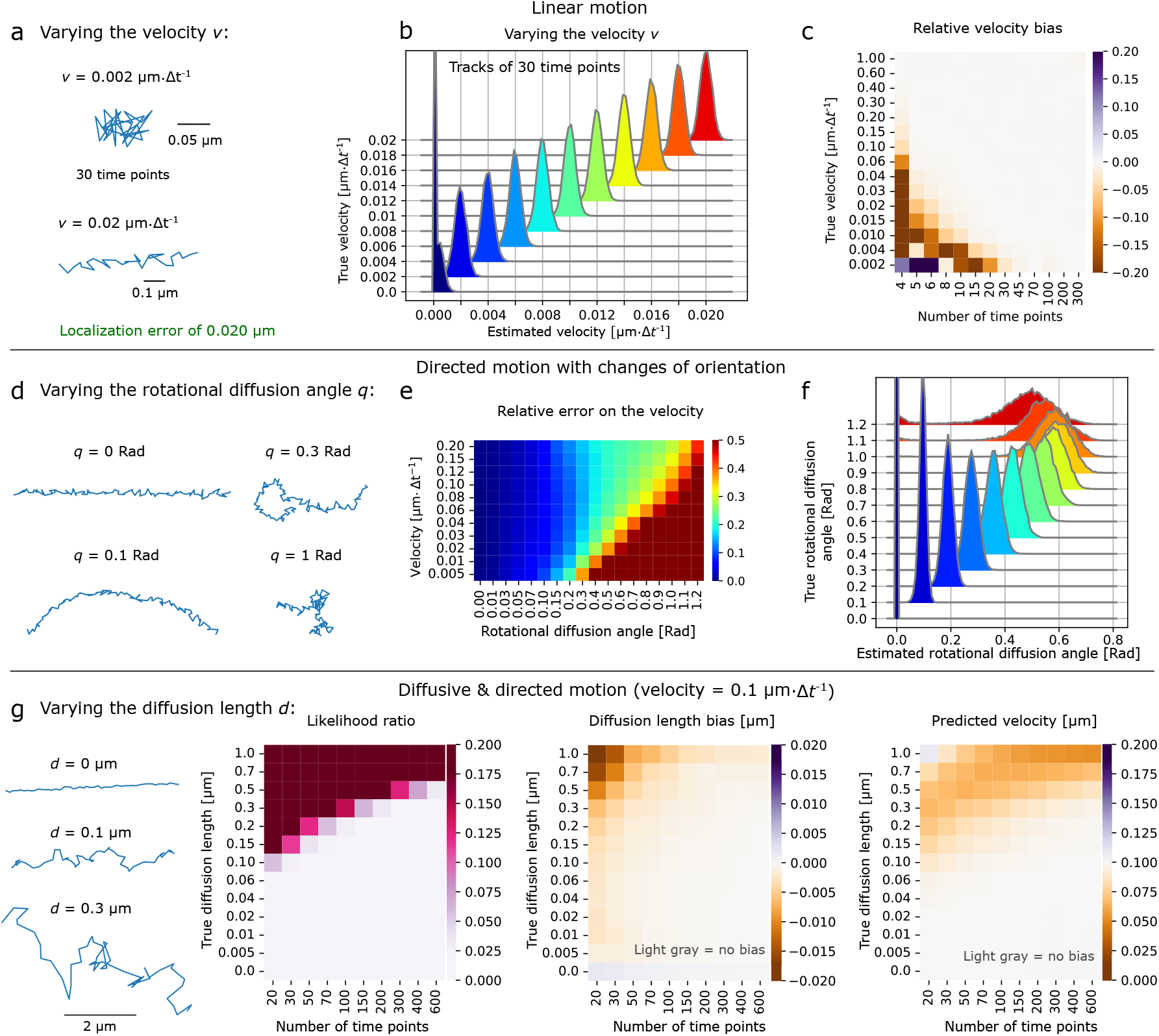
Characterizing directed motion with *aTrack*. **a-c**: Tracks with linear motion (constant speed and orientation). **a**: Examples of simulated tracks in directed motion. **b**: Histograms of the estimated velocity of individual tracks of 30 time points. 10,000 tracks per histogram. True parameters *d* = 0. µm, Localization error *σ* = 0.02 µm (fixed). The next panels use the same parameters unless specified otherwise. **c**: Heatmap of the relative biases on the estimated velocity 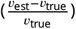. **d-f**: Tracks with constant speed but changing orientation. **d**: Simulated directed tracks with rotational diffusion. Here, the rotational diffusion angle coefficient is defined as the standard deviation of the change of orientation at each time step (analogous to the diffusion length). *v* = 0.02 µm·Δt^−1^. **e**: Heatmap of the error on the rotational diffusion angle for a range of velocities and rotational diffusion angles. Tracks of 200 time points. **f**: Distributions of the estimated rotational diffusion angle for a range of rotational diffusion angles. Tracks of 200 time points with *v* = 0.1 µm·Δt^−1^. **g**: Tracks simultaneously undergoing both linear motion and diffusion with varying levels of diffusion. Heatmaps of the likelihood ratio, bias on the diffusion length in µm, and estimated velocity depending on the number of time points per track and on the diffusion length *d*. **a-g**: Where not stated otherwise, the track parameters were as following. Localization error *σ* = 0.02 µm, *d* = 0.0 µm, velocity *v* = 0.1 µm·Δt^−1^, constant speed and orientation.

In real experiments, persistent motion is rarely perfectly linear, that is, changes in direction and speed are very common (7, 18–20). Naturally, characterizing directed motion with direction (orientation) changes is more difficult when localization error is non-negligible (21, 22). To verify our method’s capacity to accurately quantify tracks with such behaviors, we simulated tracks with constant speed and random changes of orientation (rotational diffusion). We previously observed similar behavior when analyzing the directed motion of the Rod complex in bacteria (23), motor-driven directed motion in mammalian cells (21, 24), and cell motility (25) . We varied the rate of orientation changes and the track velocity to determine the working range of aTrack for this type of directed motion (Fig. 4d,e,f). In Fig. 4e,f, we find accurate estimates of the velocity and the rotational diffusion for a wide range of parameters. As the velocity increases, we can distinguish directed motion from Brownian motion for an increasing range of rotational diffusion (Fig. 4e). However, we also see that fast changes in orientation (high rotational diffusion) make estimating the velocity and rotational diffusion more difficult. This tradeoff is expected, as rapid changes in direction make the track appear more diffusive as shown in the likelihood ratio heatmap Fig. S5a. The diffusion length heatmap (Fig. S5a) explains why the rotational diffusion coefficient is poorly estimated when high: the model interprets the high rotational diffusion as simple diffusion since the two types of motion are very difficult to distinguish in this regime.

Sometimes, particles undergo diffusion and directed motion simultaneously, for example, particles diffusing in a flowing medium (26). Sample drift can also introduce a combination of diffusion and directed motion in single-molecule tracking. Indeed, the thermal expansion of instrument components like microscope stages can induce steady motion that masks the biologically relevant diffusive motion (27). We first tested that our method could correctly differentiate directed motion from diffusion (Fig. 4g) for a range of diffusion coefficients and track lengths at a fixed velocity of 0.1 µm.Δ*t*^−1^. For mixed motion, our method accurately estimates the diffusion length and velocities even for short tracks, provided the diffusion length was low compared to the velocity. When the diffusion length is large compared to the velocity, parameters can still be predicted, but longer tracks are needed for reliable estimates.

#### Characterizing populations of tracks

The amount of information in an individual track is limited by its length making it difficult to extract the parameters precisely. One way to overcome this limitation is to consider a population of tracks that have the same state. The likelihood of the population can be easily computed by multiplying the probabilities of the individual tracks. To test our population approach, we simulated two populations: confined tracks and directed tracks. Fig. S6 shows that the expected linear increase in the log likelihood with the number of tracks and that using more tracks results in more precise parameter estimates.

Populations of tracks can often contain multiple states (e.g., free diffusion and directed motion). By taking into account the fraction of the particle in each state, we built a multi-state population model. We tested the capacity of our multistate-population model on groups of simulated tracks with 300 time points, where each track follows one of the 5 states shown in Fig. 5a.

**Figure 5.**
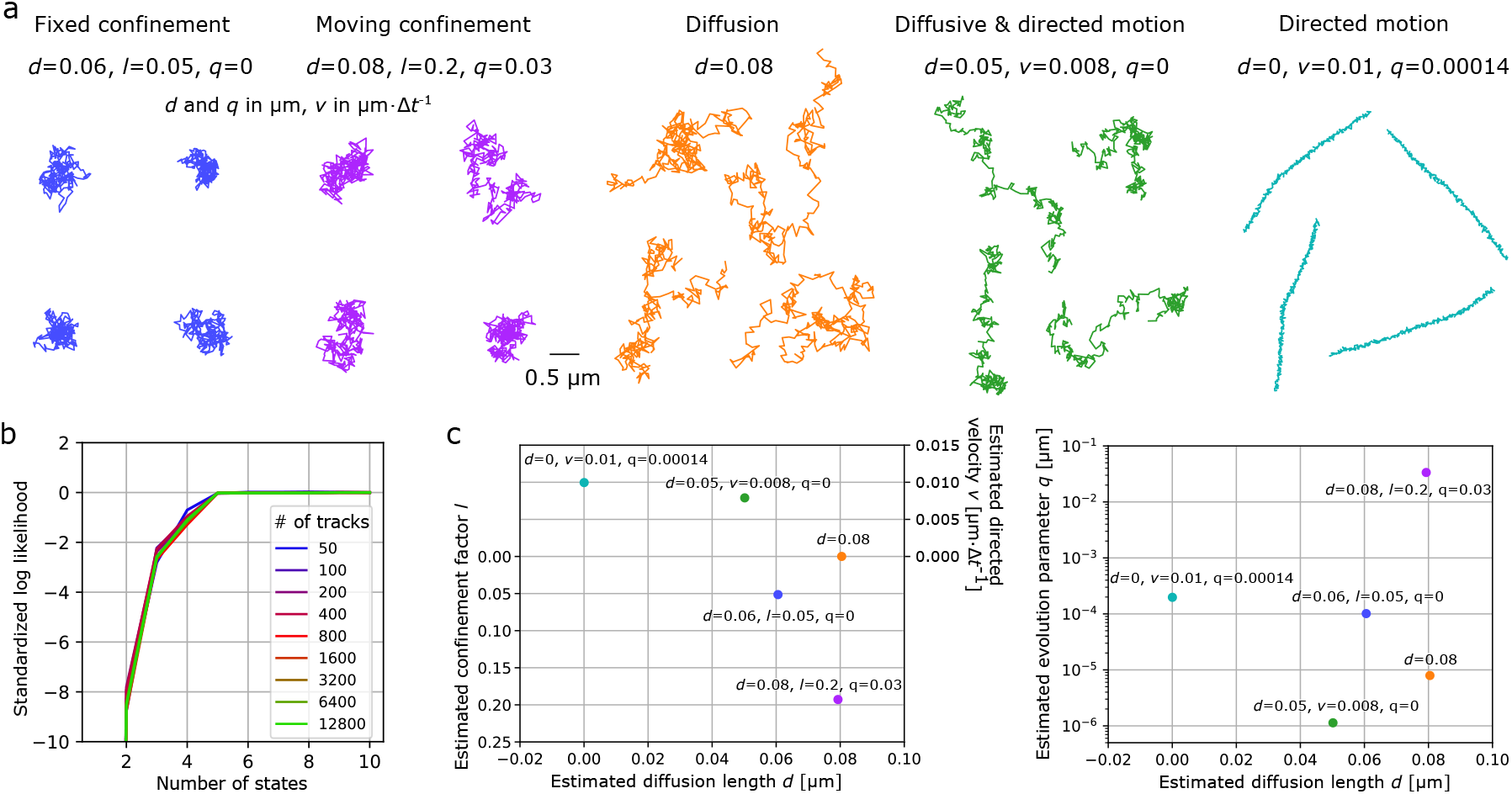
Characterizing populations of multiple states. Analysis of tracks with 5 sub-populations of set diffusion length *d*, confinement factor *l*, velocity *v* for directed tracks, and anomalous change parameter *q* (diffusion length of the potential well for confined tracks and changes of speed for directed tracks). Tracks are 300 time point long. **a**: Track examples from each of the 5 states with the corresponding state parameters. **b**: Log likelihood of the model depending on the number of states assumed by the model. The log likelihood was normalized by the number of tracks, offset by the log likelihood assuming 10 states. **c**: Estimated parameters for the 5 states (using a 5-state model).

The first step is to determine the number of states. To this end, we compute the likelihood of the model depending on the number of states. As expected, the likelihood increases with the number of states, until the correct number of states is reached and the function plateaus, in this case 5 states (Fig. 5b). This plateau in likelihood is a good indicator that the appropriate number of states is reached; however, increasing the number of states further usually results in higher likelihood, albeit marginally.

Quantitative criteria such as the Akaike Information Criterion (AIC) (28) and the Bayesian Information Criterion are often used to determine the number of parameters of a model by placing a trade-off between increasing the likelihood and increasing the number of parameters. When increasing the number of tracks, from 50 to 12,800, we found these criteria to be unreliable (Fig. S7). In theory, these criteria only work if the true underlying model is included in the alternative models, which is never the case for real tracks. As an alternative, we found that adding a small penalization term proportional to the number of parameters and the log likelihood provides a reliable criterion for identifying the number of states for any dataset size, even with mismatches between the data and the model assumptions.

Once we have identified the number of states, the parameters of each state are estimated at the population level (Fig. 5b), even when individual tracks remain difficult to classify. For instance, classifying whether a track is in the diffusive state (orange) or diffusive plus directed state (green) is difficult.

### Robustness to model mismatches

One of the most important features of a method is its robustness to deviations from its assumptions. Indeed, experimental tracking data will inevitably not match the model assumptions to some degree, and models need to be resilient to these small deviations. To test the generalizability of our approach to other types of motion, we simulated tracks using a different motion model, namely fractional Brownian motion (29). This was performed with three anomalous diffusion exponents, 0.5, 1.0, and 1.5, corresponding to sub-diffusion, Brownian diffusion, and super-diffusion, respectively Fig. 6a. The performance of aTrack was then measured by computing the difference between the likelihood of the directed-motion model and the confined-motion model. This metric is closely related to the likelihood ratio mentioned earlier, and has the advantage of showing all three motion behaviors. We expect the likelihood difference to be <0 for sub-diffusive motion, near 0 for Brownian diffusion, and >0 for super-diffusion. We verified this by plotting a histogram of the log-likelihood differences for each type of motion and estimated the classification accuracy in detecting anomalous diffusion from Brownian diffusion. aTrack was 88% accurate when differentiating subdiffusive fractional Brownian motion (anomalous diffusion exponent of *α* = 0.5) from Brownian motion (*α* = 1), and 95% accurate for differentiating superdiffusive fractional Brownian motion (*α* = 1.5) from Brownian motion.

**Figure 6.**
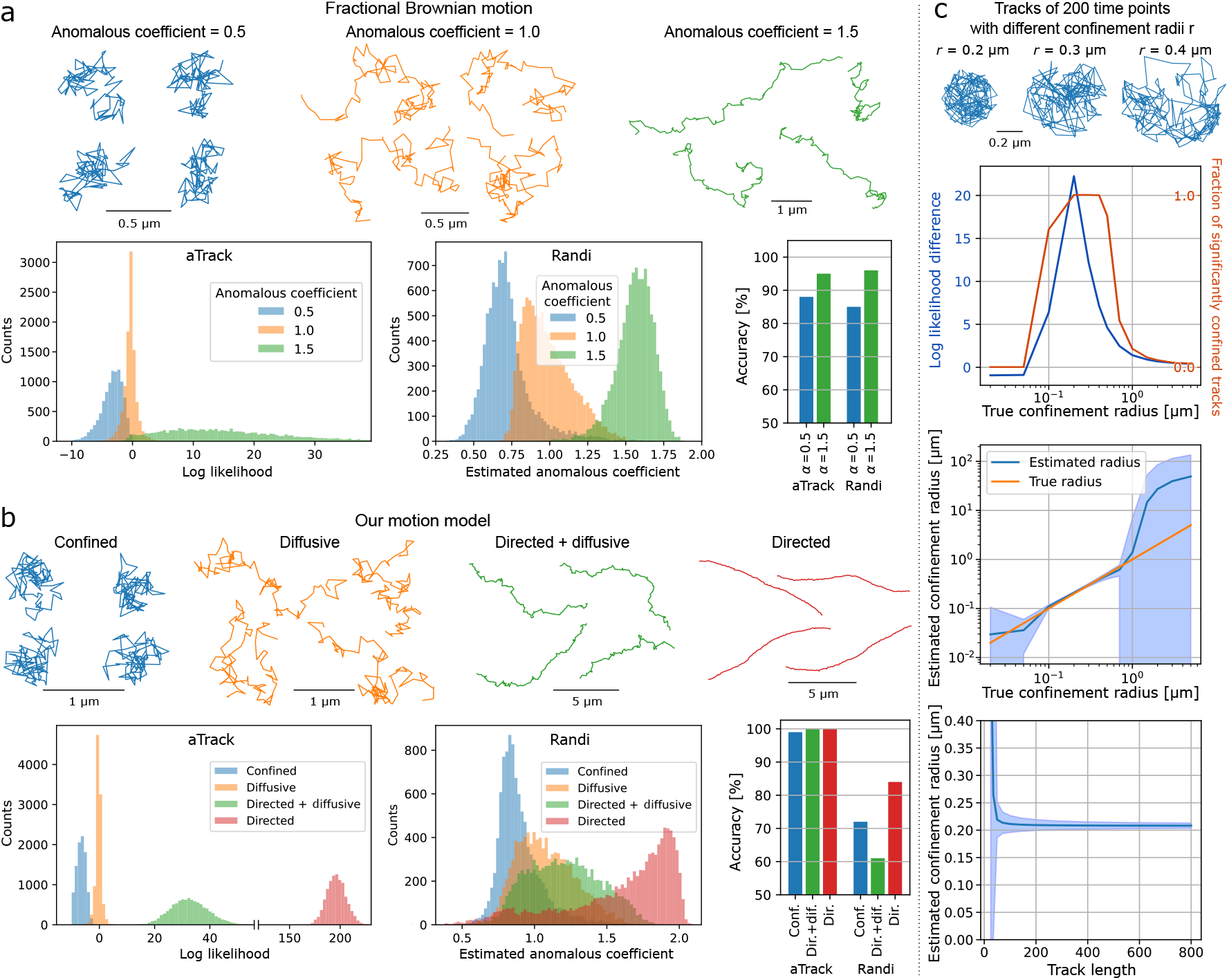
Model robustness with other motion types. **a-b**: Example tracks and corresponding distributions used to determine the type of motion for aTrack and Randi (30). aTrack uses the difference between the likelihood assuming super-diffusion and the likelihood assuming sub-diffusion (bottom-left). To classify tracks using Randi, we used the estimated anomalous exponent. The accuracy is the fraction of correctly labeled tracks in a data set composed of 5,000 sub-diffusive or superdiffusive tracks and 5,000 Brownian tracks. Classifications were done using the thresholds that best divide the distributions. **a**: Analysis of tracks with 200 time steps following fractional Brownian motion with anomalous exponent of 0.5 (subdiffusive), 1 (diffusive), and 1.5 (superdiffusive). **b**: Analysis of tracks with 200 time steps following our motion model. Confined tracks: diffusion length *d* = 0.1 µm, localization error *σ* = 0.02 µm, confinement force *l* = 0.2, fixed potential well. Brownian tracks: *d* = 0.1 µm, *σ* = 0.02 µm. tracks in both directed and diffusive motion: *d* = 0.1 µm, *σ* = 0.02 µm, directional velocity *v* = 0.1 µm·Δt^−1^. Directed tracks: *d* = 0. µm, *σ* = 0.02 µm, *v* = 0.1 µm·Δt^−1^, angular diffusion coefficient 0.1 Rad^2^·s^−1^. **c**: Analyzing tacks confined by hard boundaries using aTrack. A simulated track with 200 time points diffusing on disks of different sizes. Top panel: Log likelihood difference *L*_*c*_ − *L*_*B*_ and fraction of significantly confined tracks (likelihood ratio *l*_*B*_/*l*_*c*_ < 0.05) depending on the confinement radius. Middle panel: Estimated confinement radius 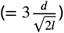 depending on the true confinement radius. Bottom: estimated confinement radius depending on the track length. Blue areas: standard deviations of the estimates.

To compare our approach to one of the leading methods specifically designed to characterize fractional Brownian motion, we performed the same test with Randi (30), the best-performing machine learning method in a head-to-head comparison of available techniques (31). On the same fractional Brownian motion dataset, Randi and aTrack achieved similar accuracies.

We then used the same approach to compare aTrack and Randi on tracks generated with our motion model, which differs from Randi’s anomalous motion models (Fig. 6b). Here, we created a dataset of anomalous diffusion with parameters that result in tracks with qualitatively similar properties and MSD curves to those observed with fractional Brownian motion (see tracks in Fig. 6a&b). For these data, our model performs with high accuracy, at least 99%. In contrast, Randi shows a lower classification accuracy. In particular, Randi has difficulty differentiating diffusive tracks from tracks with both diffusion and directed motion, 61% accuracy (only 11% better than random labeling). Curiously, directed versus Brownian tracks were also surprisingly inaccurate (83%), considering the striking difference in track behaviors (see Fig. 6b, diffusive versus directed examples).

To test another type of mismatch between our hidden variable model assumptions, we simulated diffusive tracks confined within rigid boundaries. This differs from our model, which uses a potential well to model confinement. We varied the radius of the rigid boundary for simulated tracks and measured the effect on the estimated confinement radius (Fig. 6c). aTrack accurately determines the confinement radius for a wide range of confinement radii, where the calculated confinement radius is estimated as 3-times the standard deviation of the potential well. The lower radius bound of the operating range relates to the diffusion length per step, while the upper bound is limited by the extent of the particle’s exploration of the confinement area. Note that the exploration distance is determined by the track length and by the diffusion length). This calculation was relatively independent of track length (Fig. 6d).

Motion blur is another experimental effect in single-particle tracking that can bias parameter estimation. While our model does not explicitly account for it, the estimated diffusion coefficient can be easily corrected to adjust for that effect (Fig.S8). As described by Berglund, static and dynamic localization errors have antagonistic effects on the offset term of the MSD. Our model, which explicitly models static localization error but not dynamic error, yields good estimates of the diffusion length if 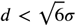 (MSD curve with a positive offset), but it underestimates *d* by a factor 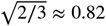 if 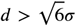. To explicitly adapt our tool to motion blur, one can include motion blur using our new framework for model design (33).

### Implementation on experimental data

To test the usefulness of aTrack on experimental data, we performed classification, population characterization, and parameter-estimation experiments.

First, we applied aTrack to analyze the movement of the spindle pole body (SPB) in Saccharomyces cerevisiae (Fig. 7a). The SPB is the microtubule-organizing center in yeast and is embedded in the nuclear envelope. Directed motion of the SPB occurs during S phase, where actin is required to establish spindle orientation by directing the spindle pole body towards the bud neck as well as during mitosis when the spindle elongates to separate the chromosome masses of mother and daughter cell (34). To visualize SPB dynamics, we imaged unsynchronized cells expressing Spc42-mCherry at a time resolution of 100 ms and used treatment with Latrunculin A (Lat A) to disrupt actin polymerization. Computing the MSD of the population of tracks showed that in average tracks appear diffusive (Fig S9a). To go beyond this ensemble metric, we used aTrack to compute the likelihood ratio (*l*_*b*_/*l*_*d*_ ) of each track of 99 time points and to infer the associated p-value from the ratio and the distribution of the likelihood ratio under the null hypothesis (Fig. 7b). Then, we computed the fractions of significantly directed tracks (type I error rate of 5%) (Fig. 7c). In untreated cells, 17 % of the tracks exhibited significant directed motion. In contrast, cells treated with LatA showed significantly lower fractions of directed tracks ( 10%, p-value = 0.0108). This drop in the fraction of directed tracks was not observed in a LatA-resistant strain. Thus, aTrack can reliably detect directed actin-dependent motion at timescales of 10 s.

**Figure 7.**
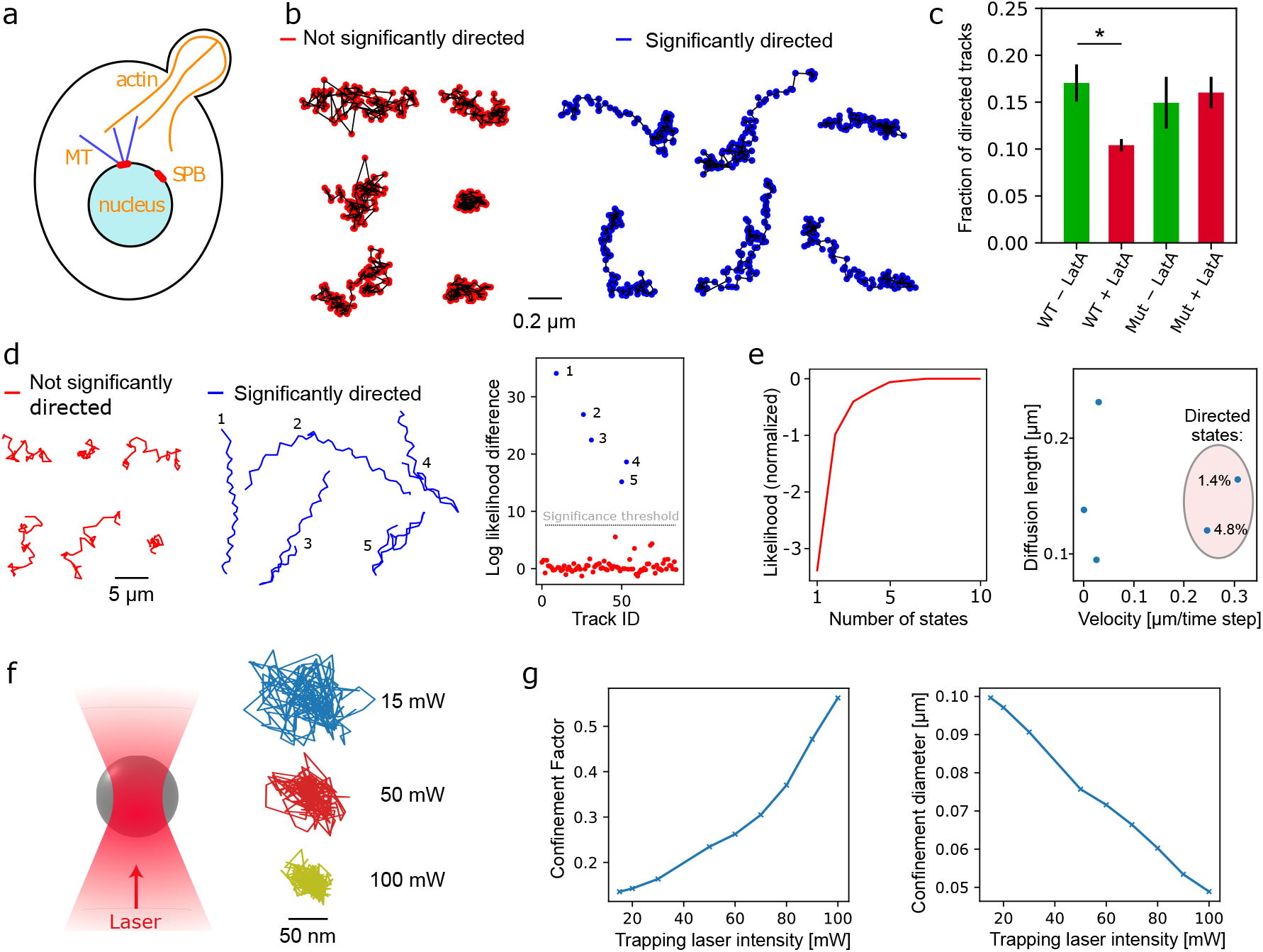
Experimental demonstrations. **a**: Illustration showing the interaction of the budding yeast spindle pole body (SPB) with actin via microtubules (MT). Actin-dependent motors are responsible for moving the nucleus toward the bud neck during S-phase. **b-c**: Analysis of spindle pole body (SPB) tracks. The analysis was carried out on tracks of 99 time points. **b**: Examples of tracks classified by aTrack to be either significantly directed or non-significantly directed. A random selections of tracks colored by their associated likelihood ratios with and without LatA treatment can be found in Fig S9b. **c**: Mean fraction of directed tracks from 3 biological replicates for the WT and 2 biological replicates for the latrunculin resistant mutant. Each replicate contains at least 682 tracks. Error bars: standard deviation. *: significant difference with a type I error of 5% according to a two-sided t-test. **d-e**: Analysis of gold nanoparticle (NP) tracks in the presence of motile bacteria (50 time points per track), where some NPs adhere to cells. **d**: NP tracks colored according to their state of motion classification using aTrack’s single-track statistical test and the log likelihood difference (*L*_*d*_ − *L*_*b*_) of all tracks. Tracks are considered significantly directed if the likelihood ratio (which is an overestimate of the p-value) is lower than 0.05 (type I error) divided by the number of tracks (85) according to the Bonferroni correction (= log likelihood difference > 7.44). **e**: Maximum likelihood (per track) of the population of tracks depending on the number of states (minus the likelihood assuming 10 states). **f-g**: Analysis of tracks for 1 µm beads trapped using optical tweezers with different laser powers. **f**: Illustration of the optical trap and example tracks of 100 time points for different laser powers. **g**: Fitting a single-state confined diffusion model on a population of 300 tracks with 20 time points.

To test the applicability of aTrack to the field of biosensing, we performed a tracking experiment with highly visible gold nanoparticles (AuNP) and *E. coli*. Specifically engineered gold nanoparticles (AuNPs) can attach to cells, and many species are motile. Zapata-Farfan et al.. Thus, detecting a directed fraction of AuNPs could be used for sensitive and fast biodetection in complex environments. AuNPs were added to a diluted *E. coli* culture and imaged. While free AuNPs diffuse rapidly, our method identified directed tracks in a 5.9% population with very high certainty (likelihood ratio < 0.001, Fig. 7d). This directed fraction is readily apparent by manual data inspection (see supplemental video). Notably, while the directed fraction also exhibited lateral oscillations, consistent with cell rotation (36), our model was robust to this deviation from the model assumptions. Next, we analyzed the tracks at the population level and computed the number of states as a function of likelihood (Fig. 7e). The likelihood function shows a noticeable difference between 4 and 5 states. In the 5-state model, there are 2 directed states and 3 diffusive states. The diffusive states likely represent free AuNPs as well as NPs conjugated to diffusing debris of different sizes. The 2 directed populations are likely caused by the abrupt tumbling motion present in some tracks. These directed states comprised 6.2% of tracks, similar to the analysis of individual tracks.

Notably, in this dataset, the likelihood plot shows an ambiguous number of states, where the likelihood increases marginally after 5 states. Such ambiguities are expected when the dataset and the model assumptions do not perfectly match. To ensure that the directed fraction is well estimated independently of the number of states, we varied the number of states to see how the parameter estimates changed, as in (37). We found that using 3-7 states, resulted in the same fraction of clearly directed tracks with similar parameters.

Finally, we tested our method’s capability to detect confinement. To do so, we confined 1-micron beads in an optical trap (38), and varied the laser power to control the trap stiffness. Based on populations of 300 tracks of 20 time points, we measured the confinement factor *l* and confirmed that it increases with the laser power of the optical trap, while the calculated confinement diameter *u* decreases, where 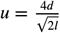 (Fig. 7g).

## Discussion

aTrack is a new tool for classifying and characterizing noisy tracks, which performs well in a diverse range of conditions. Specifically, the framework’s flexibility is relevant for a wide variety of diffusive, confined, and directed motion types. Our tool classifies the motion type as Brownian or not and quantifies biologically relevant motion parameters, such as the diffusion coefficient, confinement diameter, and velocities. Importantly, this approach calculates how statistically robust a classification is regarding the likelihood difference from a Brownian diffusion model. The flexibility of our approach in capturing a variety of motion-type behaviors contains the usefulness of the catch-all approach of using an anomalous exponent; however, unlike the anomalous exponent that fits multiple underlying motion models (6), our framework has the advantage of outputting interpretable parameters that describe the motion, e.g. velocity or confinement radii.

aTrack has several features that make it advantageous for analyzing directed and confined motion. For example, allowing the anomalous variable to change over time enables more flexibility. For directed tracks in cells, motion is rarely exactly straight over long distances. Allowing changes in velocity makes it possible to classify and characterize these curved trajectories robustly. Analogously for confined motion, a confined particle may leave its local environment or hop between environments (39–41). Our model accounts for this by allowing the center of the confinement well to diffuse as well. Interestingly, we found that directed motion can be readily identified from very short tracks due to the deterministic nature of this type of motion. Longer tracks are needed to classify confined motion with the same statistical certainty. This is to be expected, as a confined particle should reach the boundary of the confinement area several times to be distinguishable from a freely diffusive particle.

As with all classification tools, a source of ambiguity arises when motion types resemble one another. For example, this confusion occurs between immobile particles and very tightly confined particles, which can sometimes be indistinguishable and not necessarily insightful. Indeed, immobilized fluorophores can be modeled as confined to an area around a substrate to which they are bound. To better distinguish these ambiguous behaviors, experimental modifications are needed, such as adapting the experimental framerate or improving the localization precision using with brighter fluorophores (42, 43).

While machine-learning tools have proven to be effective for subdiffusion characterization (31), our approach shows it is possible to achieve similarly high performance with many fewer parameters. For our tool, these parameters map onto the stochastic physical behaviors of particle motion, making interpretation more straightforward. Compared to the machine-learning tool we tested, we found that aTrack is more robust to small model mismatches. This finding is consistent with the well-documented issues of machine-learning models generalizing poorly to new data. Of course, in the context of tracking, the generalizability of a model to new data is a key factor, as experimentally-obtained data never perfectly match the model assumptions or the training data set. One solution to make a machine learning model that generalizes better is to use a physics-informed neural network (44). Such a network would use probabilistic relationships to efficiently learn the physical properties of unlabeled tracks and would contain far fewer parameters than classical networks.

It is often useful to assume a finite number of states with fixed parameters to model the various molecular states in a sample (43). Selecting the right number of states is difficult, but can be automated by different methods. The criterion, e.g., AIC and BIC, used to determine the number of states is reliable when the model and the data are in perfect agreement. However, experimental data never match the model assumptions perfectly, and even discrepancies between models and simulated data can affect reliability, such as continuous-time simulations and discrete models. This usually results in overestimating the number of states for large data sets (45). To avoid this flaw of classical criteria, we showed that a penalization term proportional to the number of states and to the log likelihood of the data can prevent the spurious increase of the number of states when increasing the number of tracks. The drawback of this approach is the addition of a tunable parameter that influences the number of estimated parameters of the model, and it may still be necessary to limit the number of states to the biologically relevant system, or consider groups of states, e.g. all significantly directed states.

An important limitation of our approach is that it presumes that a given track follows a unique underlying model with fixed parameters. In biological systems, particles often transition from one motion type to another; for example, a diffusive particle can bind to a static substrate or molecular motor (46). In such cases, or in cases of significant mislinkings, our model is not suitable. However, this limitation can be alleviated by implicitly allowing state transitions with a hidden Markov Model (15) or alternatives such as change-point approaches (30, 47, 48), and spatial approaches (49).

In conclusion, we have shown that aTrack can identify anomalous diffusion and parameterize the motion over a broad range of motion types using a robust probabilistic framework. As the motion-model parameters estimated by the method represent physical phenomena, these variables are readily interpretable, for example the diffusion coefficient, confinement radius, and velocities. Finally, the employed motion models were selected to permit analytical integration, which makes calculating the model parameters fast and accurate; of course, this integration strategy can be implemented for a variety of motion models with hidden states, further expanding the applicability of this approach to other motion types.

## Methods

### Modeling particle motion with observed and hidden variables

#### Probabilistic model for confined motion

At each time step *i*, our confined-motion model consists of (1) a Brownian motion step, (2) a confinement step, (3) an update to parameters, and (4) a localization error step. The Brownian motion step updates the particle’s position *r*_*i*_ to an intermediate position *z*_*i*_. The variable *z*_*i*_ −*r*_*i*_ follows a Gaussian distribution centered at 0 with standard deviation *d*, where *d* is the diffusion length, 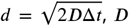 is the diffusion coefficient, and Δ*t* is the time step. Next, the confinement step is modeled by an attractive force between the particle with a potential well centered at *h*_*i*_. More precisely, the particle moves toward the center of the potential well proportionally to a confinement factor *l* and to the distance *r*_*i*_ −*h*_*i*_. The next real position *r*_*i*+1_ is thus determined by the following relationship *r*_*i*+1_ = (1 −*l*) · *z*_*i*_ +*l* ·*h*_*i*_. To allow the potential well to move, *h* is updated at each step such that *h*_*i*+1_ − *h*_*i*_ follows a Gaussian distribution of mean 0 and standard deviation *q*. Finally, to model localization error, the observed positions *c*_*i*_ and the real positions *r*_*i*_ are related by the localization precision, *σ*, where *c*_*i*_ − *r*_*i*_ follows a Gaussian distribution of mean 0 and standard deviation *σ*.

The distribution of positions for a diffusive particle in a fixed potential well is a Gaussian distribution with standard deviation 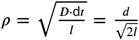. As we have no prior information about the initial center of the potential well, we assume it is positioned according to a Gaussian probability density function centered on the initial observed position. While it would be even better to consider a Gaussian 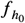 centered around *r*_0_, we simplify it by approximating it to be centered around *c*_0_ and of standard deviation 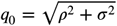. The joint probability density function corresponding to this model is the product of Gaussian functions shown in equation 1, which is integrated over all hidden parameters to calculate the likelihood, *l*_*conf ined*_ .

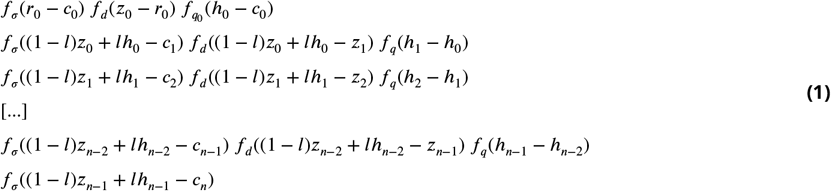

Where the real positions of the particle *r*_*i*_, potential well *h*_*i*_, and intermediate position *z*_*i*_ are hidden variables, and *l* is the confinement factor. By integrating this joint probability over the hidden variables, we can retrieve the probability of the track (the observed positions) given the model and its parameters.

While this integration step can be computed with a Monte Carlo approach (9), this is computationally expensive. Instead, we integrate using an analytical recurrence formula. This formula is allowed by the fact that the joint probability density function is a product of Gaussians and by the property that for two Gaussians, *f* and *g*, with means *μ*_*f*_, *μ*_*g*_ and standard deviations *σ*_*f*_, *σ*_*g*_, the product is also Gaussian, *f* (*x*) · *g*(*x*) = *ϕ* · *η*(*x*) where *ϕ* and *η* are Gaussian distributions described by Equation 2.

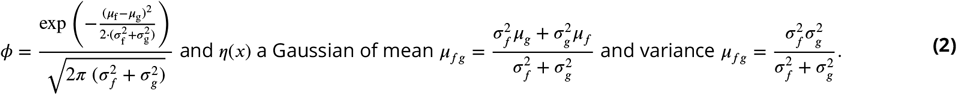

See Supplementary information for more details.

#### Probabilistic model for directed motion

To consider directed motion, we use the same general framework as confinement. At each time step *i*, the real particle position *r*_*i*_ is first updated to an intermediate position *z*_*i*_, where the variable *z*_*i*_ −*r*_*i*_ follows a Gaussian distribution centered at 0 with standard deviation *d*. Next, we add the directed-motion component with a vector *w*_*i*_, such that the next real position is the sum of the two steps, *r*_*i*+1_ = *z*_*i*_ + *w*_*i*_. Combining these two substeps, we get *z*_*i*_ − *r*_*i*_ = *r*_*i*+1_ − *r*_*i*_ − *w*_*i*_, simplifying the integration process of the probability density function expressed in equation 3. Analogous to our confinement model, the velocity vector, *w*_*i*_, is allowed to change over time, where the *w*_*i*+1_ −*w*_*i*_ follows a Gaussian distribution with mean 0 and standard deviation *q*. Finally, we include the effect of localization error, where the observed position *c*_*i*_ is related to the real position *r*_*i*_ following a Gaussian distribution, where *c*_*i*_ −*r*_*i*_ is Gaussian distributed with mean 0 and standard deviation *σ*.

During the first time step, the orientation and length of the directed motion vector (speed) need to be initialized. To do so, we assume *w*_0_ follows a Gaussian distribution function with mean 0 and standard deviation *v*. The resulting joint probability density function is shown in Equation 3.

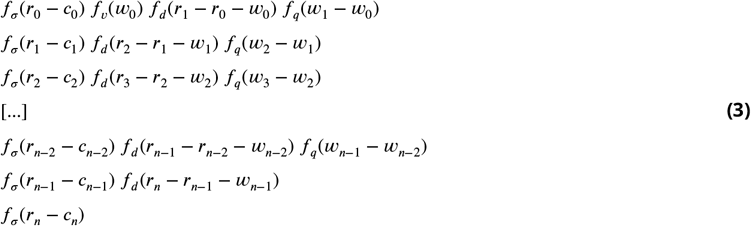

As with confinement, the probability of a directed track given the model parameters can be calculated using an analytical recurrence formula to integrate over the hidden positions, *r*_0→*n*_, and velocities *w*_0→*n*−1_. The parameter *v* can, in principle, be used to estimate the velocity of a particle with a constant speed but changing orientation in the imaging plane; however, to estimate the average speed of a particle, we use another metric, *k*, that appears in our integration process (see Supplementary information).

#### Modeling time-varying velocities and changes in direction

In our model, each axis, x and y, is treated separately, and the velocities can evolve according to a Gaussian distribution. This allows a particle’s direction to change over time. To quantify how direction changes affect the analysis for particles with a constant speed, we simulated tracks with a fixed speed and time-dependent direction for a range of angular diffusion coefficients, *D*_*θ*_. The model parameter, *q*, represents the standard deviation of the change in speed, which can be converted to an angular diffusion length using the following trigonometric relation and the estimated speed of the motion.

In the case of pure directed motion with direction changes, let us consider a single time step, *i*. We have A, the previous particle position *r*_*i*−1_; B, the particle position assuming no changes of orientation or diffusion, *r*_*i*−1_ + *v*_*i*−1_; and C, the actual particle position after a change of orientation, *r*_*i*_. ABC forms an isosceles triangle, which can be split into 2 right triangles. We find the following relationship between the scalar distances BC, AB, and 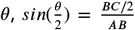, where *θ* is the orientation angle change.

#### Fitting method

For the pure Brownian model, the parameters are the diffusion coefficient and the localization error. For the confinement model, the parameters are the diffusion coefficient, the localization error, confinement factor, and the diffusion coefficient of the potential well. For the directed model, the parameters are the diffusion coefficient, the localization error, the initial velocity and the acceleration variance.

These parameters are estimated using the maximum likelihood approach which consists in finding the parameters that maximize the likelihood. We realize this fitting step using gradient descent via a TensorFlow model. All the estimates presented in this article are obtained from a single set of initial parameters to demonstrate that the convergence capacity of aTrack is robust to the initial parameter values.

#### Multi-state population model

We designed a specific algorithm to retrieve the number of states in a data set and estimate the parameters of each state as described in the code of the script *atrack*.*py* available on our Github page https://github.com/FrancoisSimon/aTrack. This multi-state population algorithm starts by performing individual fittings of tracks to get a type of motion and a set of parameters for each track. Then, we use a Gaussian mixture model on the parameters of the individual tracks to provide an overestimate of the number of states (e.g. 20 if we expect 5 states) and fit the model so that every actual state underlying the data is well represented by at least one of the the model state. Next, we iteratively remove the least useful model state and refit the model until we obtain a single-state model. At each iteration, the least useful state is determined as the state with the smallest negative impact on the likelihood.

#### Track simulations

We subdivided each time step into 20 substeps to simulate approximately continuous tracks. We applied the Brownian diffusion and anomalous movements to each of these substeps. For confined motion, the latter step moves the particle toward the center of the potential well center proportionally to the distance multiplied by a scaled confinement factor, *l*/20. In simulations with a moving potential well, the center moves according to the well’s diffusion coefficient, *q*. For directed motion, we applied a shift of constant velocity at each time step after the diffusion step. The orientation of the directed motion could vary according to a rotational diffusion coefficient. The particle’s position after the 20 substeps was set to the particle’s real position, *r*_*i*_, and the localization error was added to create the observed position *c*_*i*_. This process is equivalent for undivided frames for directed motion, as the diffusion and the directed motion steps are independent. However, for the confined model, where the diffusion of the particle influences the anomalous step, thus there is a dependence between the diffusion and confinement steps.

Fractional Brownian motion was simulated using the Python package ‘fbm’ (https://pypi.org/project/fbm/) with the Davies and Harte method (50).

### Experimental methods

#### Spindle-pole body

Yeast cells expressing Spc42-mCherry with or without allelic mutation in actin act1-113 (51), (strains KWY10722 and KWY10328 described in (52)) were grown to exponential growth phase in synthetic complete medium with glucose (SCD). Cells were treated with 0.2 mM Latrunculin A (Enzo Life Sciences, BML-T119-0500) and DMSO as solvent control for 10 min prior to imaging. Matrical 384-well glass bottom plates coated with Concanavalin A were used to image treated cells on a temperature-controlled inverted Nipkow spinning disk microscope equipped with the Yokogawa Confocal Scanner Unit CSU-W1-T2 controlled by the VisiVIEW Software (Visitron). It was used in spinning disk mode with a pinhole diameter of 50 µm combined with a 1.45 NA, 100x objective. Images were acquired on an EMCCD Andor iXon Ultra camera (1024x1024 pixel, 13x13um pixel size); for one of three biological replicates, the data was acquired with dual camera settings. Imaging was performed at 30 °C with 80 % laser intensity of a Diode 561 nm, 200 mW laser. Timelapse data were acquired with 100 ms exposure time in stream mode for 300 frames. For all movies, tracks were obtained using the ImageJ (53) plugin TrackMate (54). Peak detection: LoG with radius = 0.45 µm and threshold = 20, linkage: simple lap tracker with a linking maxiumum distance of 0.5 µm and allowing gaps of one frame. The fractions of directed tracks were inferred by computing the likelihood ratio for each track of 99 time points. Then, we simulated Brownian tracks of diffusion length 0.1 µm which corresponds to the diffusion length of the most diffusive tracks (99 percentile) to estimate the distribution of the likelihood ratio of tracks following the null hypothesis. This is a conservative estimate as we found that higher diffusion lengths (compared to the localization error) produce likelihood ratio distributions that are less skewed toward 1. Based on these simulations, we found that tracks with a likelihood ratio lower than 0.295 are significantly directed with an error rate of 5%.

#### Bacteria detection and tracking

Bacteria detection with nanoparticles was performed following the protocol described in Zapata-Farfan et al. (2023). In brief, *E. coli* (strain 25922, ATCC) were cultured for 24 h in Trypticase soy broth (TSB) at 37°C and then incubated for 30 min with 100 nm spherical gold nanoparticles at 50 µg/mL (A11-100-CIT, Nanopartz) in 1x PBS. Samples were placed in a small chamber consisting of double-sided tape (5 µm thickness, Nitto) between a coverslip and glass slide. Trackings was performed using Trackmate 7.0 (54) with the LoG detection method with a diameter of 0.70 µm (6 pixels) and quality threshold of 24.

#### Optical tweezers

The tracking of microspheres captured in an optical trap was achieved using a custom instrument constructed on a modular inverted microscope (MIM/RAMM, ASI Imaging), incorporating optical trapping and imaging paths. The trapping diode laser (SNP-06E-100, Teem) emitting at 1064nm (TEM_00_) had an average output power of 60 milliwatts. The laser beam is expanded using a 1:3 telescope (Achromatic doublets, Thorlabs) to overfill the objective’s back focal plane. This expanded beam was focused through a 100X, 1.45 NA objective (UPlanXApo 100X, Olympus) to trap dielectric particles. The same objective was used to collect bright-field illumination, which was imaged using a CMOS camera (ORCA - Flash4.0 LT3, Hamamatsu). A 3-axis piezo-driven stage (MicroScan SCXYZ100, Thorlabs) with a precision of 25 nm facilitated sample movement to capture particles in the trap. One µm polystyrene microbeads (Monodisperse fluorescent microspheres, Cromtech Research Center) were suspended in water. The solution was squeezed between two coverslips (No. 1.5, Thermo), and a single bead was brought into the laser trap by translating the sample. Data acquisition was performed at 315 frames per second for 2 minutes, these movies were then analyzed in shorter, 100-frame segments. Beads positions were tracked using TrackMate (54). aTrack analysis was performed on 300 tracks of 20 time points, inputting a fixed localization error and diffusion coefficient, which were determined from the lowest laser intensity (15mW) *σ* = 0.0116µm, *d* = 0.0259µm, respectively. Confinement radius 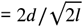.

## Data and code availability

Tracking data and the aTrack software is available online. aTrack is available as a stand-alone software for Windows and as a python package. Installation instructions are provided on the Github page https://github.com/FrancoisSimon/aTrack.

## Acknowledgements

The authors thank Sven van Teeffelen for helpful discussions. This work was supported by the Natural Sciences and Engineering Research Council of Canada [NSERC Discovery grant to MM, (RGPIN-06404-2016) to CB, (RGPIN-2022-05142) to LEW], the Canada First Research Excellence Fund (TransMedTech Institute), the ETH research grant ETH-33 19-1 and the Swiss National Science Foundation (project number 320030-236124) to ED.

## Author contributions

Conceptualization: FS, LEW; Methodology: FS, LEW; Software: FS, IF; Validation: FS; Formal analysis: FS; Investigation: FS, GR, JZ, SP; Resources: MM, CB, ED, LEW; Data curation: FS, LEW; Writing - original draft: FS, LEW; Writing - review and editing: FS, GR, SP, ED, LEW; Visualization: FS, JZ, GR, ED; Supervision: LEW; Project administration: LEW; Funding acquisition: MM, CB, ED, LEW.

## Competing interest statement

The authors declare no competing interest.

## Supplementary information

### Statistical tests

Statistical tests are important as they allow one model to be chosen over another. In the case of independent and identically distributed variables, the log likelihood ratio of a dataset drawn from the null hypothesis follows a chi-squared distribution that can be used to determine the p-value. The p-value can then be used to reject or not the null hypothesis.

### Bounding the p-value with the likelihood ratio

For a given trajectory **c** = {*c*_0_, *c*_1_, …, *c*_*n*_}, we construct likelihood ratio tests comparing a null hypothesis *H*_0_ of Brownian diffusion against an alternative hypothesis *H*_1_ of anomalous diffusion (confined or directed). The likelihood ratio is defined as 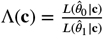, where 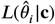 denotes the maximum likelihood under hypothesis *H*_*i*_ with maximum likelihood estimator 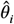.

While under standard regularity conditions, Wilks’ theorem (16) establishes that −2 log Λ(**c**) asymptotically follows a *χ*^2^ distribution with degrees of freedom equal to the difference in parameter dimensions between models, this theorem cannot be rigorously applied here as the null hypothesis assumes that the anomalous parameter lies at the boundary of its parameter space (under *H*_0_, the anomalous variable follows a Dirac delta function) (17, 55). Through simulation (Fig. S2), we empirically found that under *H*_0_, Λ(**c**) consistently concentrated near 1 (track lengths were tested from 5 to 400 time points). This shows that under these conditions, the likelihood ratio Λ(**c**) satisfies *P* (Λ(**c**) ≤ *α* | *H*_0_) ≤ *α*. As the p-value is by definition uniform and as we can define a bijection between the p-value and the skewed Λ(**c**), for a desired Type I error rate *α*, we can reject *H*_0_ when Λ(**c**) < *α*. This guarantees *P* (reject *H*_0_ | *H*_0_ true) ≤ *α*. Since the Brownian model is nested within both confined and directed motion models with confinement factor *l* = 0 or velocity *v* = 0 respectively, we have 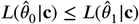 which ensures 0 ≤ Λ(**c**) ≤ 1.

### Simulation based p-value determination

While easy to implement, we acknowledge that the above method has two flaws: 1) it is based on the empirical observation that the Λ(**c**) distribution is skewed towards 1. This may not hold for tracks outside the tested range, e.g. those with more than 400 data points. 2) Using an inequality instead of realizing the bijection between Λ(**c**) and the p-value lacks power. The more Λ(**c**) under *H*_0_ concentrates near 1, the greater is this lack of power. Therefore, for applications requiring more precise p-value estimates, a simulation-based calibration procedure can be performed. Given an observed trajectory **c** with test statistic Λ_obs_, we first estimate null model parameters as 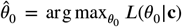. We then simulate *N* trajectories under *H*_0_ with parameters 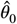 and compute the likelihood ratios 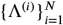. The p-value is estimated as 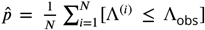. By the law of large numbers (56), 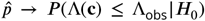 as *N* → ∞. When analyzing individual trajectories from a population of tracks, multiple testing corrections become essential to control for false positives. For *m* trajectories with individual p-values 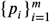, we can apply either the Bonferroni correction (57), rejecting 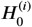 if *p*_*i*_ < *α*/*m*,or the Benjamini-Hochberg procedure (58) to control false discovery rate at level

### Multi-model comparisons

The likelihood ratio framework extends to multi-model comparisons. For model selection among *K* candidates, one can employ information criteria including the Akaike Information Criterion (28), 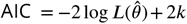, or the Bayesian Information Criterion (59), 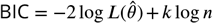, where *k* is the number of parameters and *n* is the trajectory length. Our empirical analysis (Figure S7) demonstrates that classical information criteria often overestimate the number of motion states, particularly for large datasets. This occurs due to model misspecification, as experimental data never perfectly match theoretical models (60, 61). To address this limitation, we introduce a penalization term proportional to both the number of parameters and log-likelihood: Modified 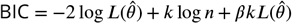, where *β* is a small positive constant that prevents spurious state proliferation.

### Derivations

#### Likelihood calculation for confined motion

To simplify the integration of the product of Gaussians described in the Method section, Eq. (1), *f*_*σ*_((1−*l*)*z*_*i*_ +*lh*_*i*_ −*c*_*i*+1_) *f*_*d*_ ((1− *l*)*z*_*i*_ + *lh*_*i*_ − *z*_*i*+1_) can be refactored to 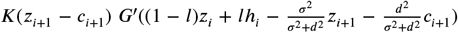 of variances *k*^2^ = *σ*^2^ + *d*^2^ and *g*^′2^ = *σ*^2^*d*^2^/(*σ*^2^ + *d*^2^). This can be further simplified into 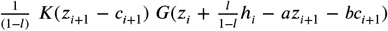 with 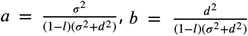 and *G* represent Gaussian distributions of mean 0 with variances *k*^2^ = *σ*^2^ + *d*^2^ and 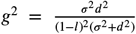, respectively. Then, we use the annotation *N*(*x, V* ) to refer to other Gaussian probability density functions of mean 0 and of variance *V* .

Eq. (1) is equivalent to:

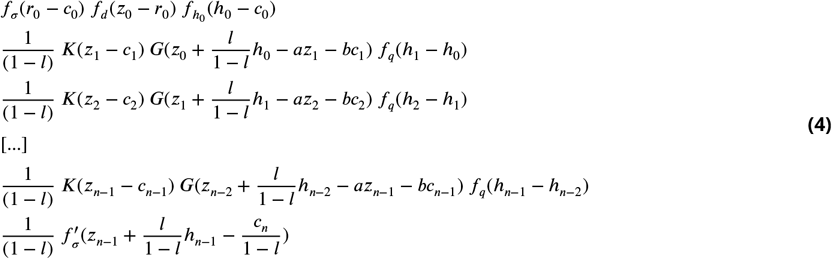

During the integration process, we obtain the following set of recurrence formulas:

- Initialization:

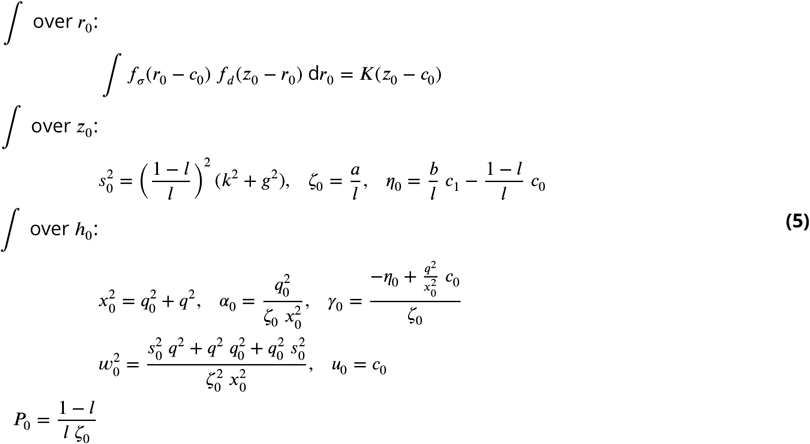
- Recurrence:

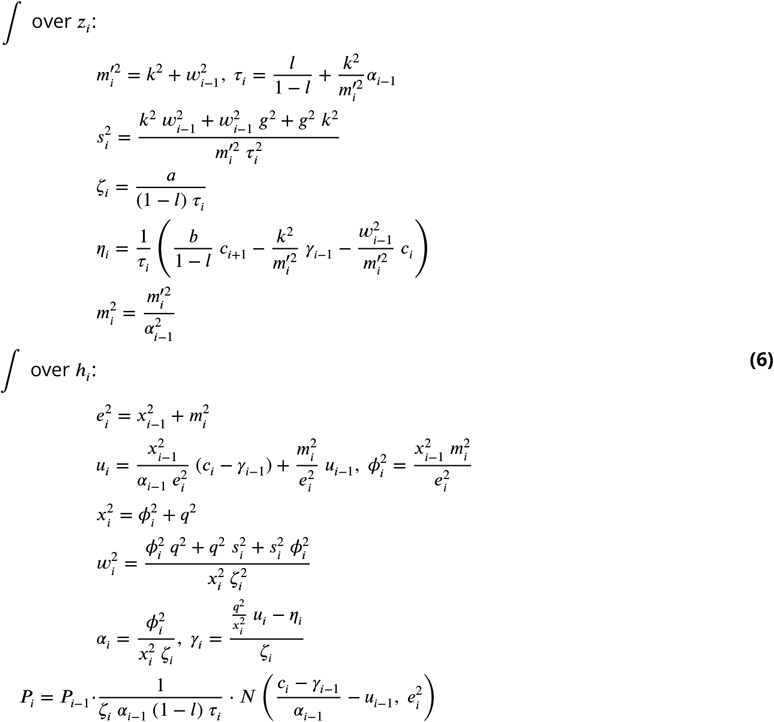
- Final step:

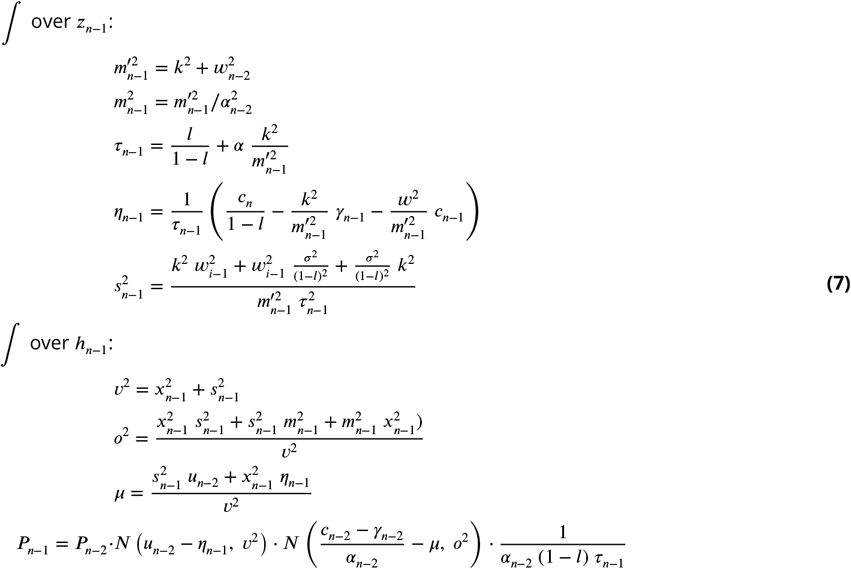

### Likelihood calculation for directed motion

Similarly to the confined motion formula, the track probability can be expressed using a recurrence formula that depends on the parameters of the directed motion model, the localization error *σ*, the diffusion length *d*, the initial velocity *v* and the speed of the velocity change *q*.

- Initialization: Step 0:

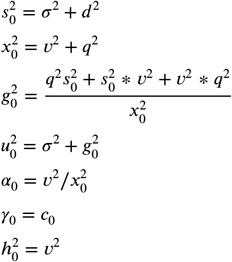

Step 1:

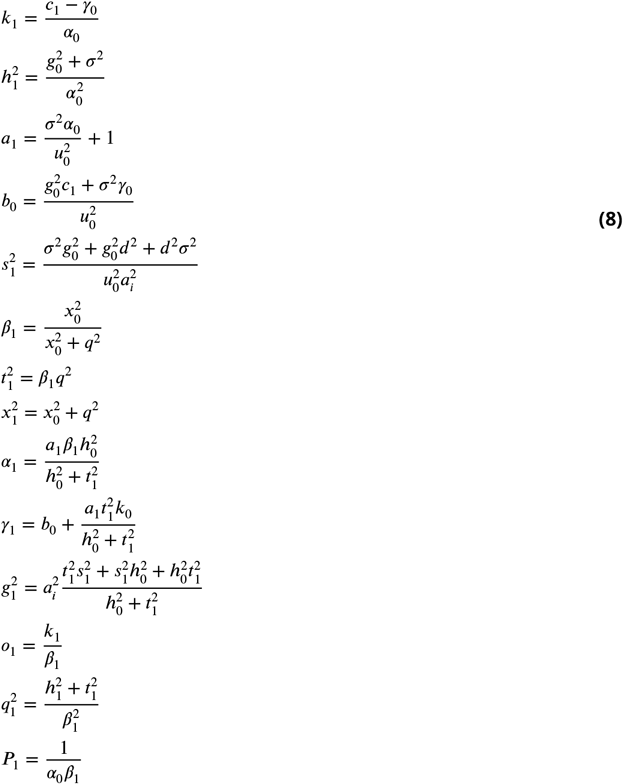
- Recurrence:

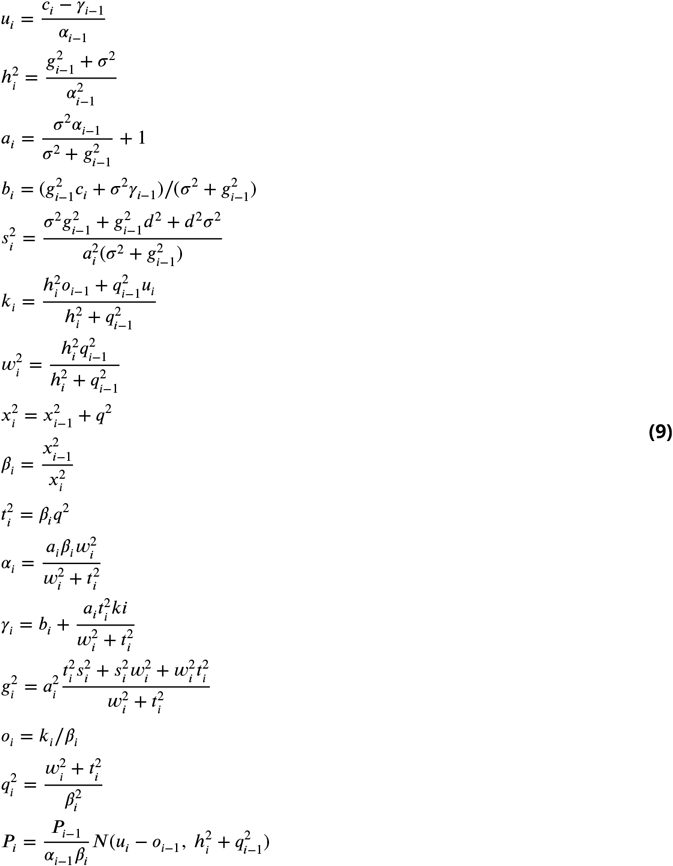
- Final steps (steps *i* = *n* − 2 and *i* = *n* − 1):

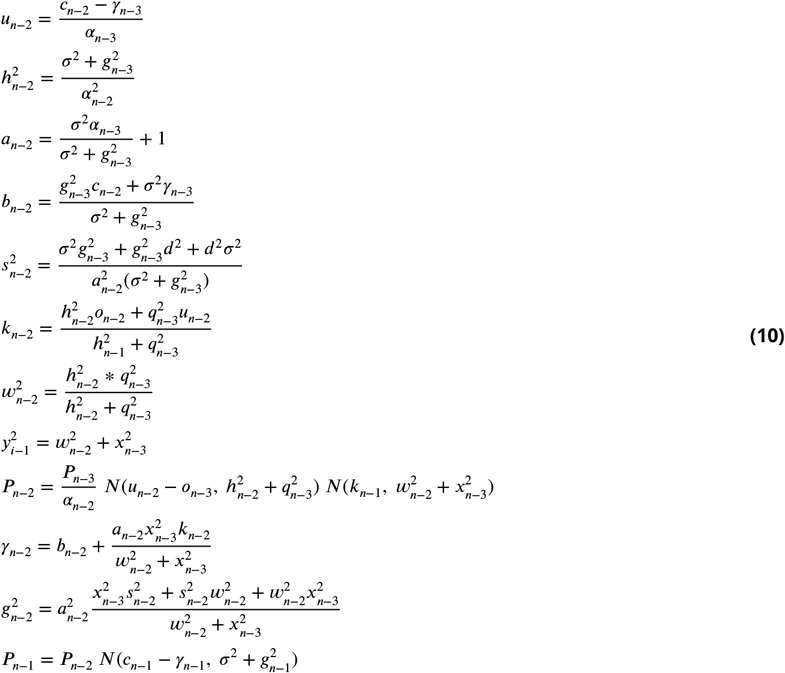

**Supplementary Figure 1.**
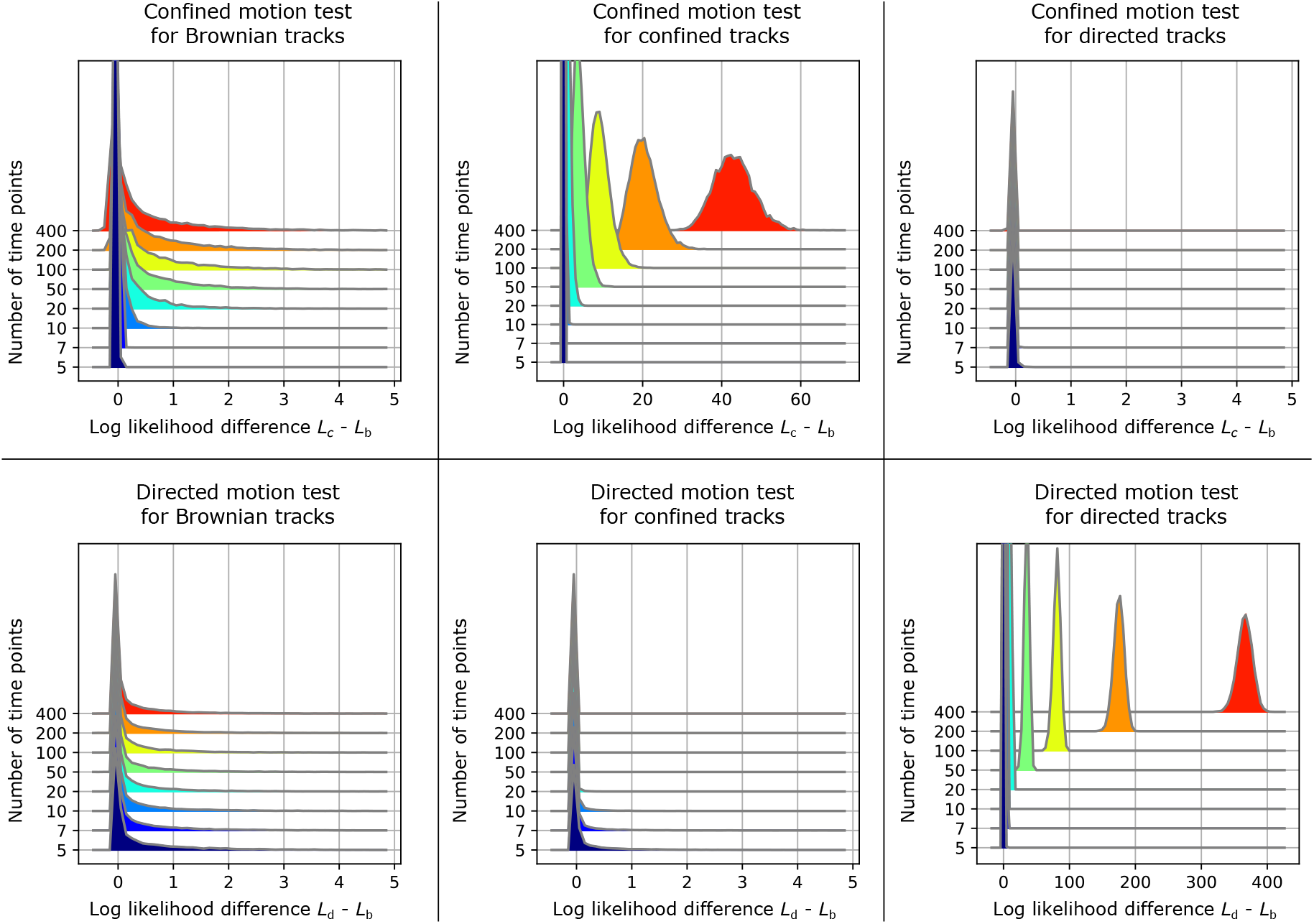
Log-likelihood differences as a function of the track length. Distributions of the log likelihood differences (*L*_*c*_ − *L*_*b*_ and *L*_*d*_ − *L*_*b*_) for tracks in Brownian motion (column 1), confined motion (column 1), or linear motion (column 3) using the confinement motion test (row 1) or the directed motion test (row 2) for tracks with different number of time points. 10,000 tracks per distribution. The simulated track parameters were as following: localization error *σ* = 0.02 µm; confined tracks: diffusion length per step *d* = 0.1 µm, confinement factor *l* = 0.25; linear tracks: *d* = 0.0 µm, velocity *v* = 0.02 µm·Δt^−1^, constant speed and orientation.

**Supplementary Figure 2.**
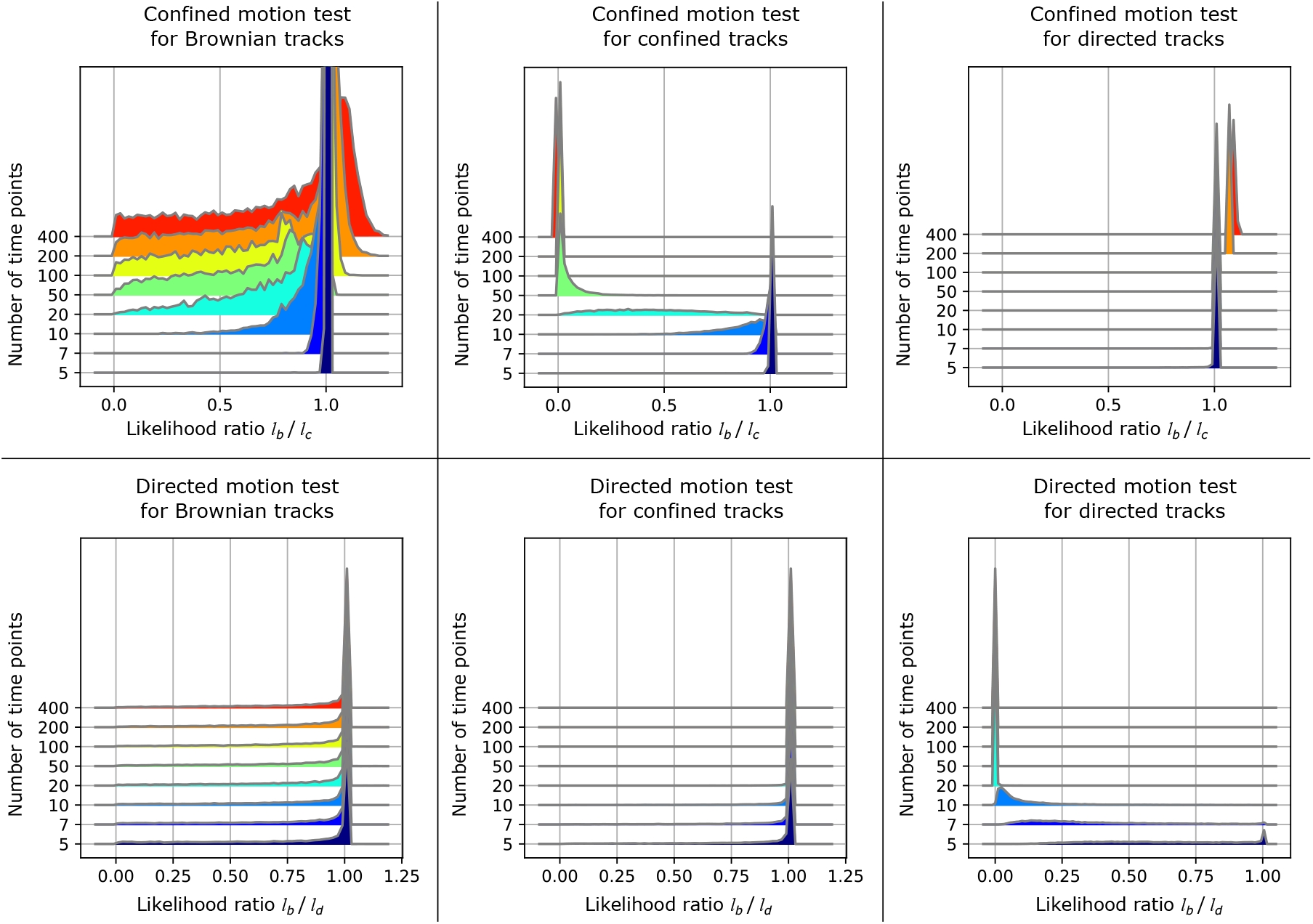
Likelihood ratios as a function of the track length. Distributions of the likelihood ratios *l*_*b*_/*l*_*c*_ and *l*_*b*_/*l*_*d*_ corresponding to Fig. S1. As expected, the distributions are skewed toward 0 only when the proper test is applied.

**Supplementary Figure 3.**
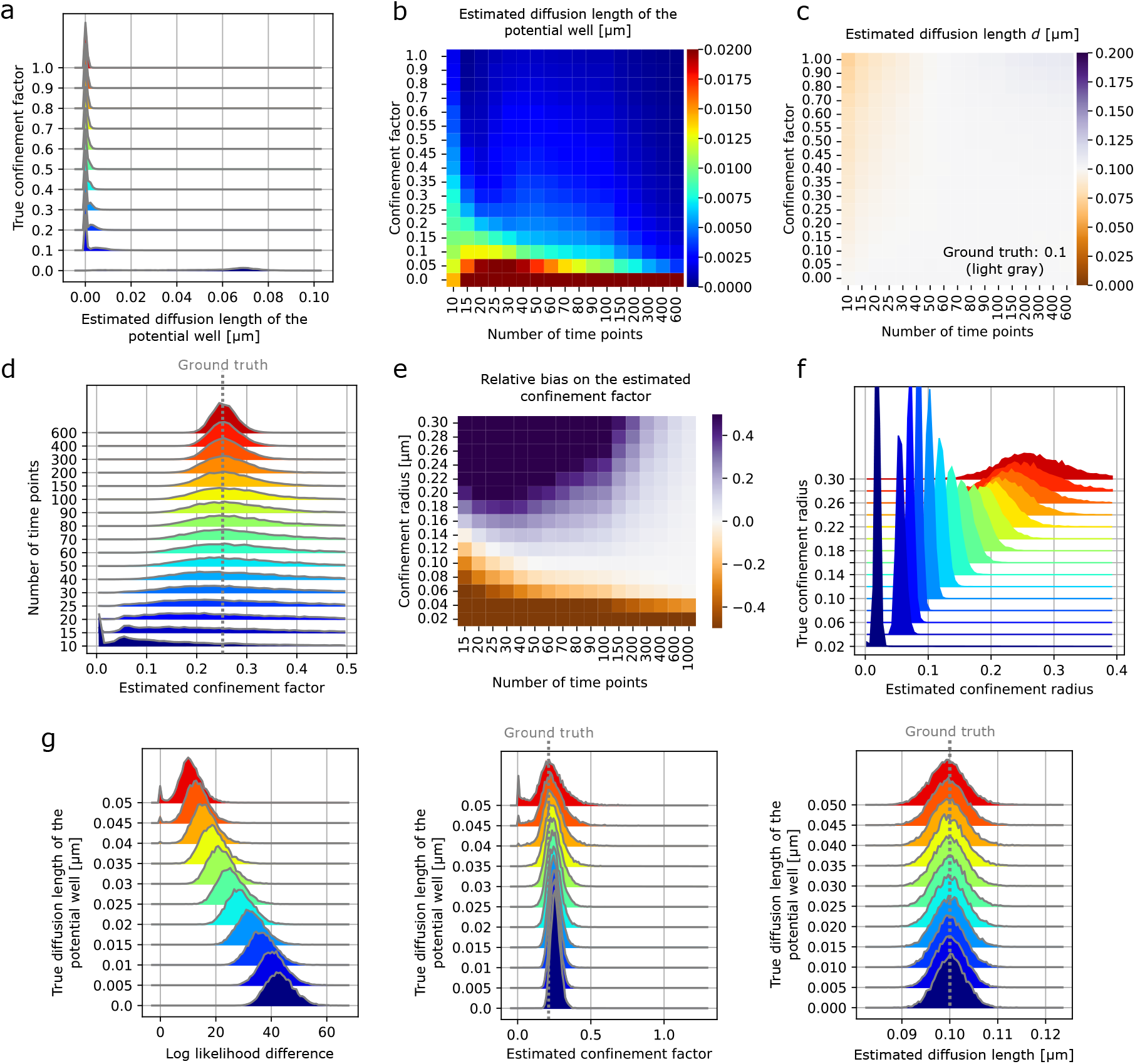
Parameter estimates for confined motion model. **a**: Histograms of the estimated diffusion length per step of the potential well depending the confinement factor corresponding to Fig 3b. **b-c**: Heatmaps of the estimated diffusion lengths per step of the potential well (b) and of the estimated diffusion lengths or the particle depending on the track length and on the confinement factor corresponding to Fig 3c. **d**: Distributions of the estimated confinement factor depending on the track length (same conditions than Fig 3c with a confinement factor of 0.25). **e**: Heatmap of the relative biases on the estimated confinement factors depending on the confinement radius and track length corresponding to Fig 3d. **f**: Distributions of the estimated confinement radius depending on the true confinement radius (same conditions as in Fig 3d with tracks of 150 time points). **g**: Distributions of the log likelihood difference (*L*_*c*_ − *L*_*b*_), estimated confinement factor and estimated diffusion length of the particle depending on the diffusion length of the potential well corresponding to Fig 3d.

**Supplementary Figure 4.**
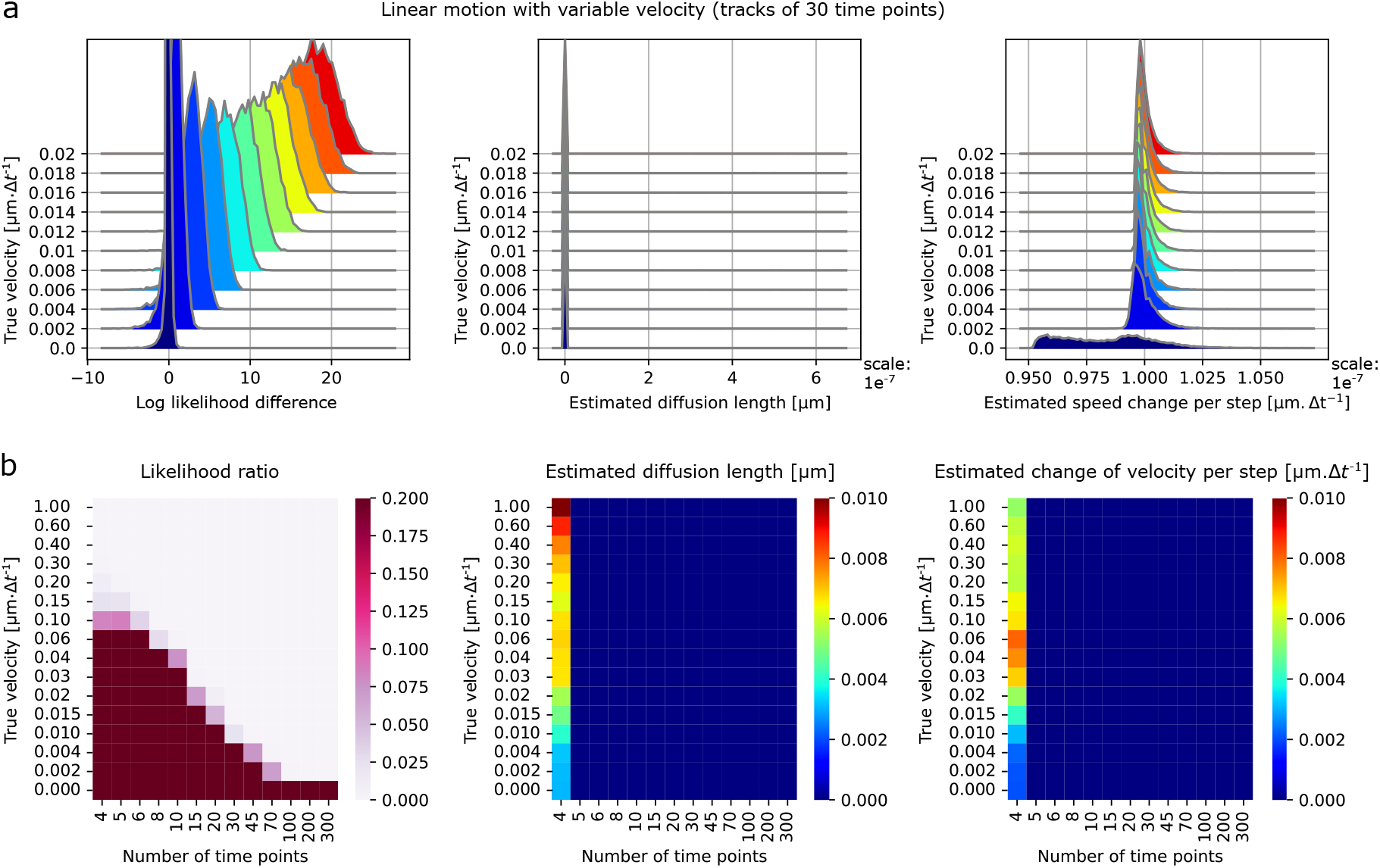
Linear motion model estimations. **a**: (Complement of Fig 4b) Rainbow plots of the log likelihood difference, estimated diffusion length, and estimated change of velocity for tracks in perfect linear motion (with localization error) for different linear motion velocities. 10,000 tracks per condition. localization error: *σ* = 0.02 µm, tracks of 30 time points. **b**: (Complement of Fig 4c) Heatmaps of the likelihood ratio, estimated diffusion length and change of velocity (average) for tracks in perfect linear motion varying the track length and the directed motion velocity. *σ* = 0.02 µm.

**Supplementary Figure 5.**
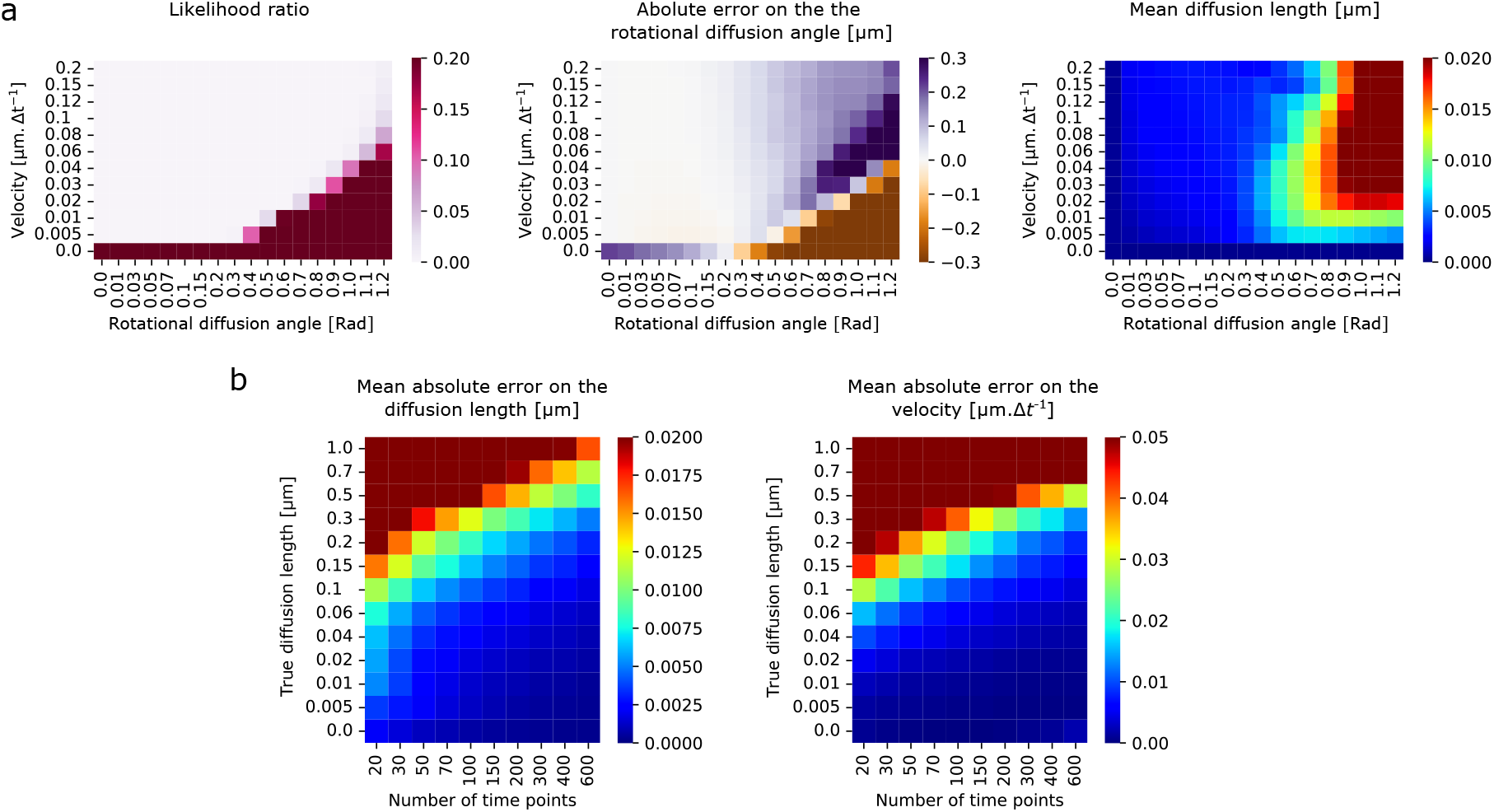
Impact of directional changes in directed motion models. **a**: (Complement of Fig 4e) Study of the impact of the rotational diffusion of tracks in directed motion with changing orientation. Heatmaps of the likelihood ratio, of the absolute error on the rotational diffusion angle, and of the estimated diffusion length when varying the directed motion velocity and the rotational diffusion angle. *d* = 0.0 µm, *σ* = 0.02 µm. **b**: (Complement of Fig 4g) Characterization of the motion parameters of particles with both diffusive and directed motion. Mean absolute error on the diffusion length and on the velocity of the linear motion varying the number of time points in each track. Directed motion velocity: *v* = 0.1 µm·Δt^−1^, *σ* = 0.02 µm. **a-b**: mean values from 10,000 tracks.

**Supplementary Figure 6.**
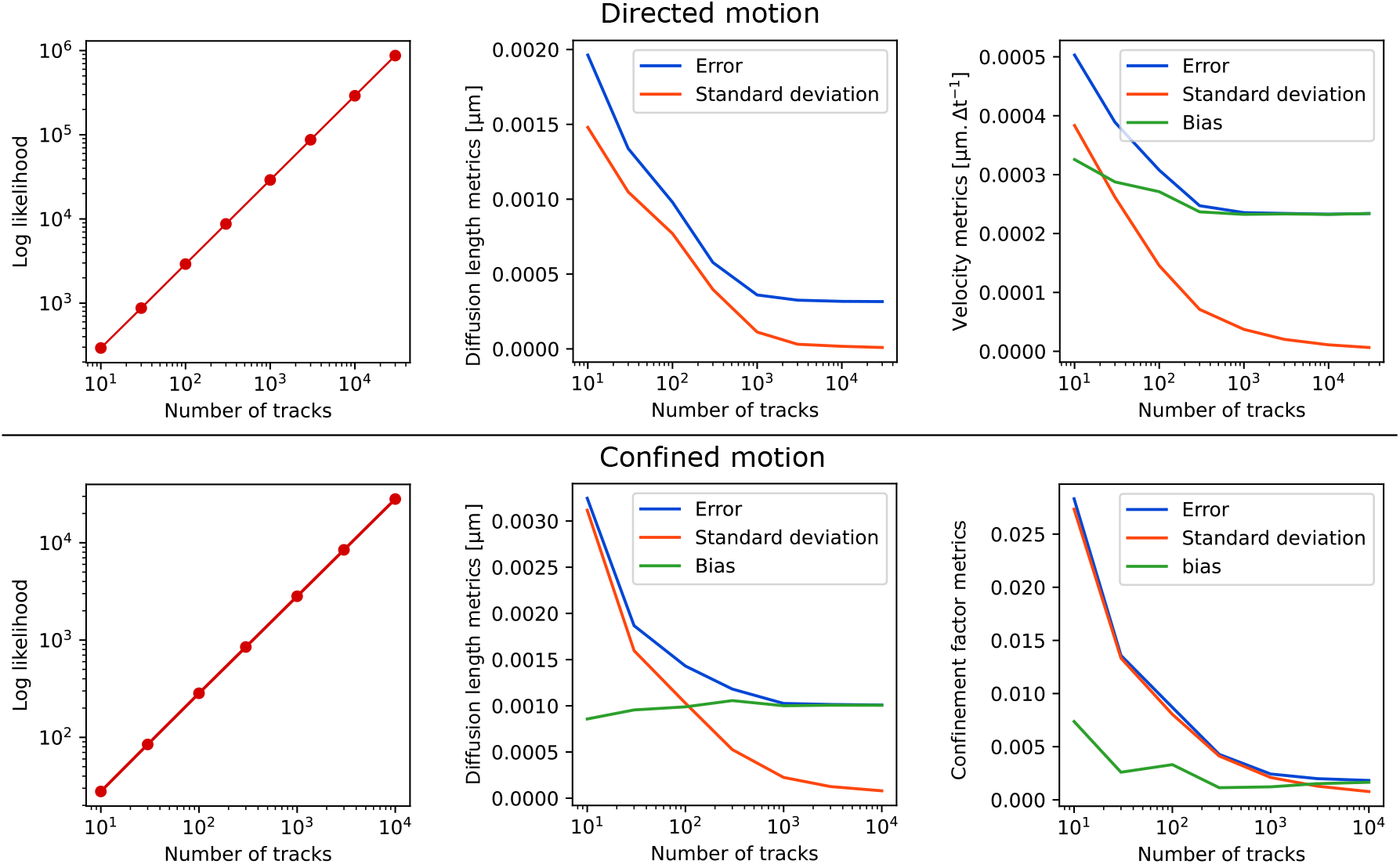
Dataset size and parameter estimate error. Effect of the number of tracks on the different parameters of the tracks: the likelihood, the root mean squared error, the standard deviation and the bias on the estimates of the diffusion length and anomalous parameter (velocity or confinement factor) for both directed motion and confined motion. All tracks were composed of 50 time points and 50 replicates were performed to estimate the error for each number of tracks. Directed tracks: persistent motion velocity *v* = 0.02 µm·Δt^−1^, angular diffusion coefficient : 0.1 Rad^2^·Δt^−1^, *d* = 0.0 µm, *σ* = 0.02 µm. Confined tracks: confinement factor 0.2, *d* = 0.1 µm, *σ* = 0.02 µm.

**Supplementary Figure 7.**
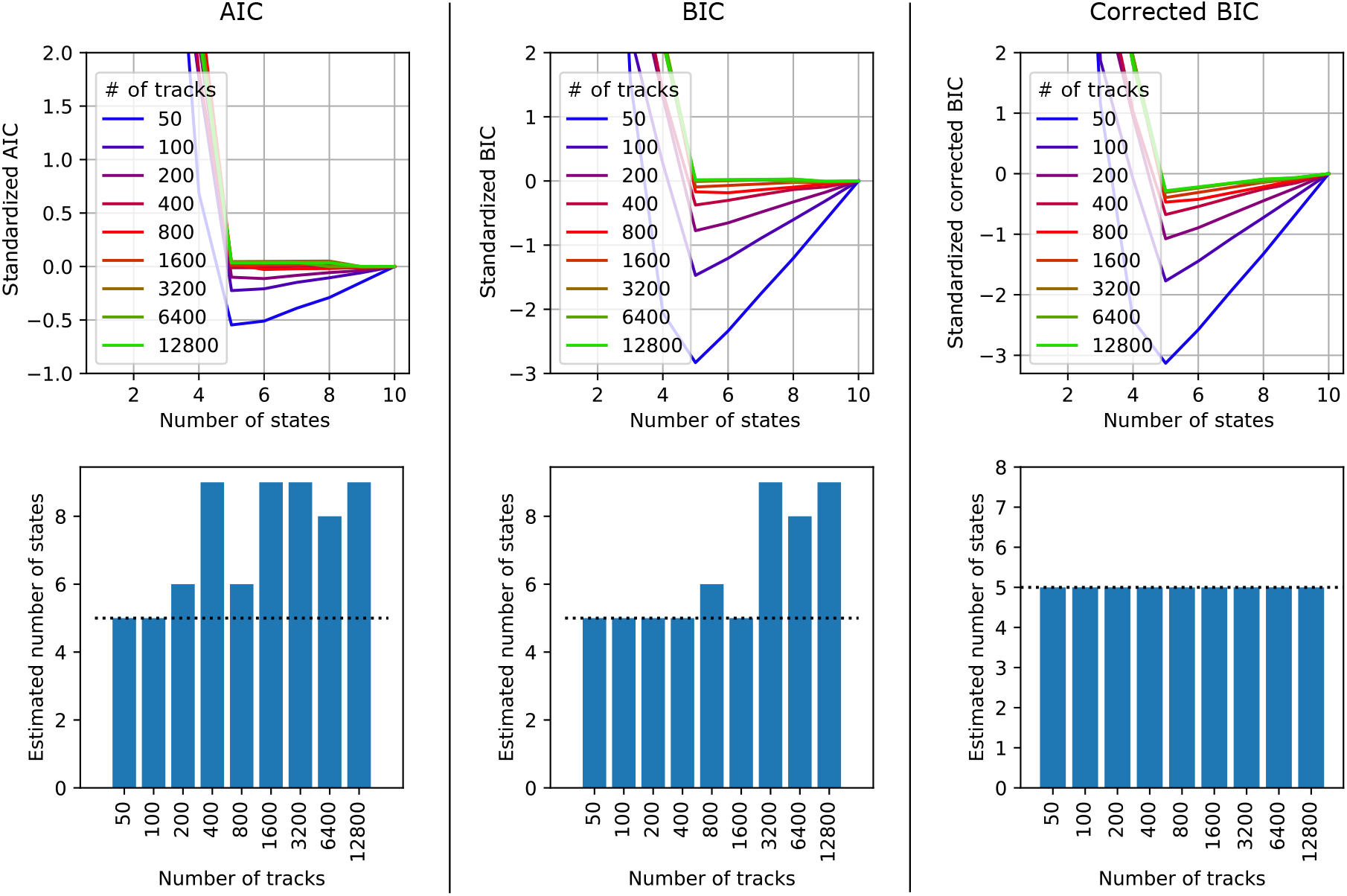
Estimating the number of states. AIC, BIC, and corrected BIC corresponding to the log likelihood shown in Fig 5c depending on the number of states assumed by the model and on the number of tracks per data set. The corrected BIC corresponds to the BIC with an additional penalization term of 0.0002*kL* with and *k* the number of parameters and *L* the log likelihood. Under the AIC, BIC, and corrected BIC curves, we plotted the optimal number of states (the one that minimizes the criterion) for each data set.

**Supplementary Figure 8.**
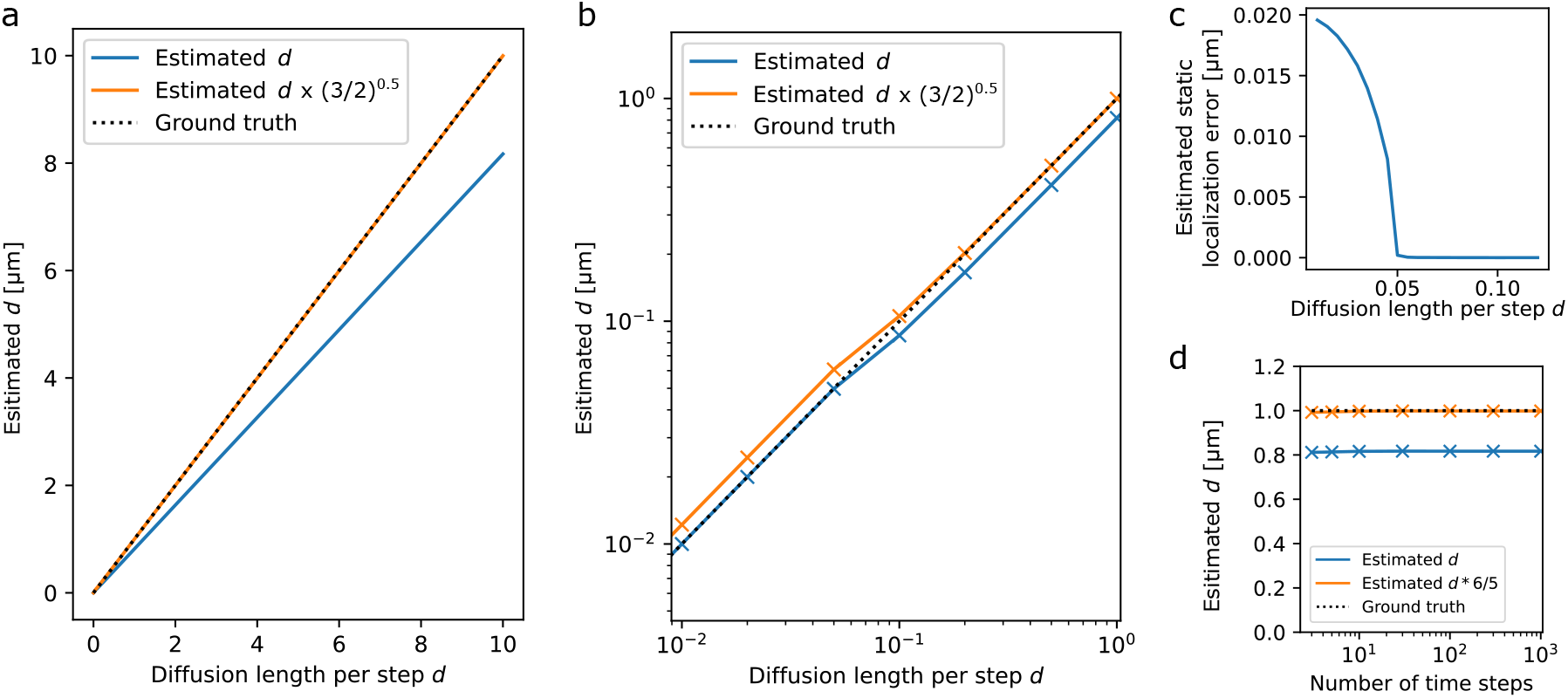
Effect of dynamic and static localization error on estimated motion parameters. Simulated populations of Brownian tracks with continuous exposure and estimated the (population-wise) diffusion length per step and localization error. At each step, the position is estimated as the average position of 200 sub-steps with a static localization error of 0.02µm (per time-step). **a-c**: Simulations of 5000 tracks of 100 time points with varying diffusion lengths. **a**: Estimated diffusion length as a function of the true diffusion length. **b**: Estimated diffusion length as a function of the true diffusion length (log-log plot). **c**: Estimated localization error as a function of the true diffusion length. **d**: Estimated diffusion lengths for 5000 tracks with varying number of time steps and fixed diffusion length per time step of 1 µm.

**Supplementary Figure 9.**
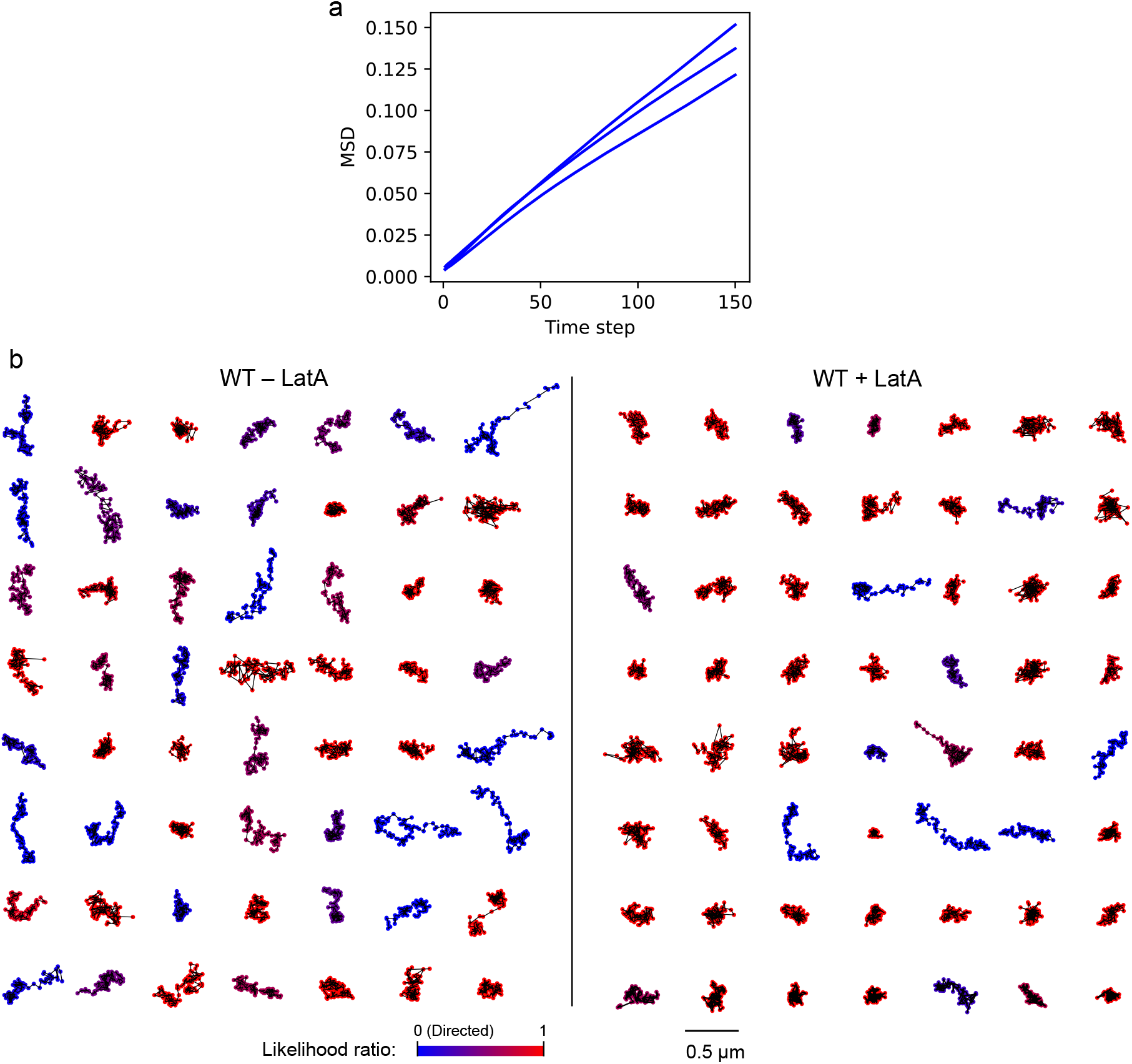
Confinement characterization in *Saccharomyces cerevisiae*. **a**: Mean squared displacements (MSD) as a function of the number of time steps in the WT strain without treatment (DMSO). **b**: Random selections of tracks of 99 time points from the WT strain without and with LatA treatment (resp. − LatA and + LatA) colored according to their likelihood ratio (*l*_*b*_/*l*_*d*_ ).

## Notes

https://github.com/FrancoisSimon/aTrack

### Competing Interest Statement

The authors have declared no competing interest.

### Summary of Updates

We have modified the supplementary section: Statistical tests

## Bibliography

1. Hao Shen, Lawrence J Tauzin, Rashad Baiyasi, Wenxiao Wang, Nicholas Moringo, Bo Shuang, and Christy F Landes. Single particle tracking: from theory to biophysical applications. Chemical reviews, 117(11):7331–7376, 2017.

2. Achillefs N Kapanidis, Stephan Uphoff, and Mathew Stracy. Understanding protein mobility in bacteria by tracking single molecules. Journal of molecular biology, 430(22):4443–4455, 2018.

3. Zhao Wang, Xuejing Wang, Ying Zhang, Weili Xu, and Xiaojun Han. Principles and applications of single particle tracking in cell research. Small, 17(11):2005133.

4. Michael J Saxton. A biological interpretation of transient anomalous subdiffusion. i. qualitative model. Biophysical journal, 92(4):1178–1191, 2007.

5. Felix Höfling and Thomas Franosch. Anomalous transport in the crowded world of biological cells. Reports on Progress in Physics, 76(4):046602, 2013.

6. Ralf Metzler, Jae-Hyung Jeon, Andrey G Cherstvy, and Eli Barkai. Anomalous diffusion models and their properties: non-stationarity, non-ergodicity, and ageing at the centenary of single particle tracking. Physical Chemistry Chemical Physics, 16(44):24128–24164, 2014.

7. Nilah Monnier, Zachary Barry, Hye Yoon Park, Kuan-Chung Su, Zachary Katz, Brian P English, Arkajit Dey, Keyao Pan, Iain M Cheeseman, Robert H Singer, et al. Inferring transient particle transport dynamics in live cells. Nature Methods, 12(9):838, 2015.

8. Jason Bernstein and John Fricks. Analysis of single particle diffusion with transient binding using particle filtering. Journal of theoretical biology, 401:109–121, 2016.

9. Paddy J Slator and Nigel J Burroughs. A hidden markov model for detecting confinement in single-particle tracking trajectories. Biophysical journal, 115(9):1741–1754, 2018.

10. Krzysztof Burnecki, Eldad Kepten, Joanna Janczura, Irena Bronshtein, Yuval Garini, and Aleksander Weron. Universal algorithm for identification of fractional brownian motion. a case of telomere subdiffusion. Biophysical journal, 103(9):1839–1847, 2012.

11. Vincent Briane, Charles Kervrann, and Myriam Vimond. Statistical analysis of particle trajectories in living cells. Physical Review E, 97(6):062121, 2018.

12. Maxime Woringer, Ignacio Izeddin, Cyril Favard, and Hugues Berry. Anomalous subdiffusion in living cells: bridging the gap between experiments and realistic models through collaborative challenges. Frontiers in Physics, 8:134, 2020.

13. Felix Oswald, Aravindan Varadarajan, Holger Lill, Erwin JG Peterman, and Yves JM Bollen. Mreb-dependent organization of the e. coli cytoplasmic membrane controls membrane protein diffusion. Biophysical journal, 110(5):1139–1149, 2016.

14. Peter K Relich, Mark J Olah, Patrick J Cutler, and Keith A Lidke. Estimation of the diffusion constant from intermittent trajectories with variable position uncertainties. Physical Review E, 93(4):042401, 2016.

15. François Simon, Jean-Yves Tinevez, and Sven van Teeffelen. Extrack characterizes transition kinetics and diffusion in noisy single-particle tracks. Journal of Cell Biology, 222(5):e202208059, 2023.

16. Samuel S Wilks. The large-sample distribution of the likelihood ratio for testing composite hypotheses. The annals of mathematical statistics, 9(1):60–62, 1938.

17. Aad W Van der Vaart. Asymptotic statistics, volume 3. Cambridge university press, 2000.

18. Paolo Pierobon, Sarra Achouri, Sébastien Courty, Alexander R Dunn, James A Spudich, Maxime Dahan, and Giovanni Cappello. Velocity, processivity, and individual steps of single myosin v molecules in live cells. Biophysical journal, 96(10):4268–4275, 2009.

19. Sven Van Teeffelen, Siyuan Wang, Leon Furchtgott, Kerwyn Casey Huang, Ned S Wingreen, Joshua W Shaevitz, and Zemer Gitai. The bacterial actin mreb rotates, and rotation depends on cell-wall assembly. Proceedings of the National Academy of Sciences, 108(38):15822–15827, 2011.

20. Aaron D Pilling, Dai Horiuchi, Curtis M Lively, and William M Saxton. Kinesin-1 and dynein are the primary motors for fast transport of mitochondria in drosophila motor axons. Molecular biology of the cell, 17(4):2057–2068, 2006.

21. Christopher P Calderon, Lucien E Weiss, and WE Moerner. Robust hypothesis tests for detecting statistical evidence of two-dimensional and three-dimensional interactions in single-molecule measurements. Physical Review E, 89(5):052705, 2014.

22. Cédric Bouzigues and Maxime Dahan. Transient directed motions of gabaa receptors in growth cones detected by a speed correlation index. Biophysical journal, 92(2):654–660, 2007.

23. Gizem Özbaykal, Eva Wollrab, Francois Simon, Antoine Vigouroux, Baptiste Cordier, Andrey Aristov, Thibault Chaze, Mariette Matondo, and Sven van Teeffelen. The transpeptidase pbp2 governs initial localization and activity of the major cell-wall synthesis machinery in e. coli. Elife, 9:e50629, 2020.

24. Stanislav Tsitkov, Yuchen Song, Juan B Rodriguez III, Yifei Zhang, and Henry Hess. Kinesin-recruiting microtubules exhibit collective gliding motion while forming motor trails. ACS nano, 14(12):16547–16557, 2020.

25. LiYan Ping. Cell orientation of swimming bacteria: from theoretical simulation to experimental evaluation. Science China Life Sciences, 55:202–209, 2012.

26. Hong Qian, Michael P Sheetz, and Elliot L Elson. Single particle tracking. analysis of diffusion and flow in two-dimensional systems. Biophysical journal, 60(4):910–921, 1991.

27. Frank J. Fazekas, Thomas R. Shaw, Sumin Kim, Ryan A. Bogucki, and Sarah L. Veatch. A mean shift algorithm for drift correction in localization microscopy. Biophysical Reports, 1(1):100008, 2021. ISSN 2667-0747.

28. Hirotugu Akaike. A new look at the statistical model identification. IEEE transactions on automatic control, 19(6):716–723, 1974.

29. Benoit B Mandelbrot and John W Van Ness. Fractional brownian motions, fractional noises and applications. SIAM review, 10(4):422–437, 1968.

30. Aykut Argun, Giovanni Volpe, and Stefano Bo. Classification, inference and segmentation of anomalous diffusion with recurrent neural networks. Journal of Physics A: Mathematical and Theoretical, 54(29):294003, 2021.

31. Gorka Muñoz-Gil, Giovanni Volpe, Miguel Angel Garcia-March, Erez Aghion, Aykut Argun, Chang Beom Hong, Tom Bland, Stefano Bo, J Alberto Conejero, Nicolás Firbas, et al. Objective comparison of methods to decode anomalous diffusion. Nature communications, 12(1):6253, 2021.

32. Andrew J Berglund. Statistics of camera-based single-particle tracking. Physical Review E, 82(1):011917, 2010.

33. Francois C Simon and Justin Cardona. Fast analytical method to integrate multivariate gaussians over hidden variables. 2024.

34. Chandra L Theesfeld, Javier E Irazoqui, Kerry Bloom, and Daniel J Lew. The role of actin in spindle orientation changes during the saccharomyces cerevisiae cell cycle. Journal of Cell Biology, 146(5):1019–1032, 1999.

35. Jennyfer Zapata-Farfan, Morteza Hasanzadeh Kafshgari, Sergiy Patskovsky, and Michel Meunier. Dynamic multispectral detection of bacteria with nanoplasmonic markers. Nanoscale, 15(7):3309–3317, 2023.

36. Thomas R. Powers. Role of body rotation in bacterial flagellar bundling. Phys. Rev. E, 65:040903, Apr 2002.

37. Joshua R Prindle, Yibo Wang, Julian M Rocha, Andreas Diepold, and Andreas Gahlmann. Distinct cytosolic complexes containing the type iii secretion system atpase resolved by three-dimensional single-molecule tracking in live yersinia enterocolitica. Microbiology Spectrum, 10 (6):e01744–22, 2022.

38. A. Ashkin and J. M. Dziedzic. Optical trapping and manipulation of viruses and bacteria. Science, 235(4795):1517–1520, 1987. doi: 10.1126/science.3547653.

39. Ido Golding and Edward C Cox. Physical nature of bacterial cytoplasm. Physical review letters, 96(9):098102, 2006.

40. Aubrey V Weigel, Blair Simon, Michael M Tamkun, and Diego Krapf. Ergodic and nonergodic processes coexist in the plasma membrane as observed by single-molecule tracking. Proceedings of the National Academy of Sciences, 108(16):6438–6443, 2011.

41. Jens Ehrig, Eugene P Petrov, and Petra Schwille. Near-critical fluctuations and cytoskeleton-assisted phase separation lead to subdiffusion in cell membranes. Biophysical journal, 100(1):80–89, 2011.

42. Keith J Mickolajczyk and William O Hancock. Kinesin processivity is determined by a kinetic race from a vulnerable one-head-bound state. Biophysical journal, 112(12):2615–2623, 2017.

43. François Simon, Lucien E Weiss, and Sven van Teeffelen. A guide to single-particle tracking. Nature Reviews Methods Primers, 4(1):66, 2024.

44. Maziar Raissi, Paris Perdikaris, and George E Karniadakis. Physics-informed neural networks: A deep learning framework for solving forward and inverse problems involving nonlinear partial differential equations. Journal of Computational physics, 378:686–707, 2019.

45. Martin Lindén and Johan Elf. Variational algorithms for analyzing noisy multistate diffusion trajectories. Biophysical journal, 115(2):276–282, 2018.

46. Lucien E Weiss, Julia F Love, Joshua Yoon, Colin J Comerci, Ljiljana Milenkovic, Tomoharu Kanie, Peter K Jackson, Tim Stearns, and Anna-Karin Gustavsson. Single-molecule imaging in the primary cilium. In Methods in Cell Biology, volume 176, pages 59–83. Elsevier, 2023.

47. Vincent Briane, Myriam Vimond, Cesar Augusto Valades-Cruz, Antoine Salomon, Christian Wunder, and Charles Kervrann. A sequential algorithm to detect diffusion switching along intracellular particle trajectories. Bioinformatics, 36(1):317–329, 2020.

48. Carlo Manzo. Extreme learning machine for the characterization of anomalous diffusion from single trajectories (andi-elm). Journal of Physics A: Mathematical and Theoretical, 54(33):334002, jul 2021.

49. Silvan Türkcan and Jean-Baptiste Masson. Bayesian decision tree for the classification of the mode of motion in single-molecule trajectories. PloS one, 8(12):e82799, 2013.

50. Robert B Davies and David S Harte. Tests for hurst effect. Biometrika, 74(1):95–101, 1987.

51. Kathryn R Ayscough, Joel Stryker, Navin Pokala, Miranda Sanders, Phil Crews, and David G Drubin. High rates of actin filament turnover in budding yeast and roles for actin in establishment and maintenance of cell polarity revealed using the actin inhibitor latrunculin-a. Journal of Cell biology, 137(2):399–416, 1997.

52. Janka Zsok, Francois Simon, Göksu Bayrak, Luljeta Isaki, Nina Kerff, Amy Wolstenholme, Lucien E Weiss, and Elisa Dultz. The nuclear basket regulates the distribution and mobility of nuclear pore complexes in budding yeast. bioRxiv, 2023.

53. Caroline A Schneider, Wayne S Rasband, and Kevin W Eliceiri. Nih image to imagej: 25 years of image analysis. Nature methods, 9(7):671–675, 2012.

54. Jean-Yves Tinevez, Nick Perry, Johannes Schindelin, Genevieve M Hoopes, Gregory D Reynolds, Emmanuel Laplantine, Sebastian Y Bednarek, Spencer L Shorte, and Kevin W Eliceiri. Trackmate: An open and extensible platform for single-particle tracking. Methods, 115:80–90, 2017.

55. Yunxiao Chen, Irini Moustaki, and Haoran Zhang. A note on likelihood ratio tests for models with latent variables. psychometrika, 85(4):996–1012, 2020.

56. Thomas S Ferguson. A course in large sample theory. Routledge, 2017.

57. Carlo Bonferroni. Teoria statistica delle classi e calcolo delle probabilita. Pubblicazioni del R istituto superiore di scienze economiche e commericiali di firenze, 8:3–62, 1936.

58. Yoav Benjamini and Yosef Hochberg. Controlling the false discovery rate: a practical and powerful approach to multiple testing. Journal of the Royal statistical society: series B (Methodological), 57(1):289–300, 1995.

59. Gideon Schwarz. Estimating the dimension of a model. The annals of statistics, pages 461–464, 1978.

60. Gerda Claeskens and Nils Lid Hjort. Model selection and model averaging. Cambridge books, 2008.

61. Kenneth P Burnham and David R Anderson. Model selection and multimodel inference: a practical information-theoretic approach. Springer, 2002.

